# Early life adversity promotes gastrointestinal dysfunction through a sex-dependent phenotypic switch in enteric glia

**DOI:** 10.1101/2024.05.31.596805

**Authors:** Jacques Gonzales, Christine Dharshika, Khadijah Mazhar, Wilmarie Morales-Soto, Jonathon L. McClain, Adam J. Moeser, Rance Nault, Theodore J. Price, Brian D. Gulbransen

## Abstract

Irritable bowel syndrome and related disorders of gut-brain interaction (DGBI) are common and exhibit a complex, poorly understood etiology that manifests as abnormal gut motility and pain. Risk factors such as biological sex, stressors during critical periods, and inflammation are thought to influence DGBI vulnerability by reprogramming gut-brain circuits, but the specific cells affected are unclear. Here, we used a model of early life stress to understand cellular mechanisms in the gut that produce DGBIs. Our findings identify enteric glia as a key cellular substrate in which stress and biological sex converge to dictate DGBI susceptibility. Enteric glia exhibit sexual dimorphism in genes and functions related to cellular communication, inflammation, and disease susceptibility. Experiencing early life stress has sex-specific effects on enteric glia that cause a phenotypic switch in male glia toward a phenotype normally observed in females. This phenotypic transformation is followed by physiological changes in the gut, mirroring those observed in DGBI in humans. These effects are mediated, in part, by alterations to glial prostaglandin and endocannabinoid signaling. Together, these data identify enteric glia as a cellular integration site through which DGBI risk factors produce changes in gut physiology and suggest that manipulating glial signaling may represent an attractive target for sex-specific therapeutic strategies in DGBIs.

## INTRODUCTION

Irritable bowel syndrome (IBS) is the most common disorder of gut-brain interaction (DGBI) and affects roughly 1 out of every 4 adults^1–3^. The etiology of IBS and related DGBIs is complex and involves neural, immune, and microbial changes along the brain-gut axis that manifest as abnormal gut motility and abdominal pain^2,4^. Bi-directional gut-brain signaling engages a vicious cycle whereby symptoms of dysmotility and pain promote, and are in turn, amplified by anxiety and stress^5^. Treatments for IBS remain unsatisfactory because the underlying biology remains incompletely understood.

Vulnerability to IBS is dictated by programming of gut-brain circuits by certain risk factors such as biological sex, exposure to stressors during critical periods in development, and inflammation. These mechanisms produce IBS in women at a rate of 1.5 to 3 times that of men, potentially due to more robust immunological responses in females^5,6^. Curiously, there are certain circumstances in which males become susceptible to developing IBS. One such example is early life stress. Over 75% of IBS patients report experiencing adverse early life experiences, and the frequency and severity of these stressful events increase the odds of developing IBS^7–9^. These observations suggest that many instances of IBS can be traced to events in early life. Males are particularly sensitive to maternal neglect compared to other types of early life adversity and exhibit greater impairments in resilience and reduced brain connections supporting autonomic regulation in IBS^9,10^. Low resilience negatively affects symptom severity and mental health in men but why males become susceptible to IBS following early life adversity is not understood. Identifying factors that drive this unique switch in sex-dependent susceptibility to IBS could ultimately benefit IBS treatment in both sexes.

Clues to the mechanisms that link the negative effects of early life adversity in males to IBS suggest a heightened cortisol response to a visceral stressor occurs mainly in men and not women^11^. In the gut, enteric glia are targets of cortisol and sympathetic neurons, and act as central regulators of neurons that control visceral sensitivity and motility^12–18^. The development of enteric glia and the neural circuits in which they reside coincides with critical periods affected by maternal separation stress^19–22^. These are times during which the developing enteric nervous system (ENS) is highly plastic and vulnerable to perturbations that reprogram gut neurocircuits that control motility and pain^5^. Enteric glia exhibit structural changes in animals subjected to maternal separation stress^23,24^, which could indicate that enteric glia are a cellular substrate that links IBS susceptibility factors such as biological sex, stress, and inflammation to dictate sex-specific changes in gut functions.

Our objective in this study was to test the hypothesis that early life stress reprograms enteric glial phenotype in a sex-dependent manner to regulate DBGI susceptibility and that these glial changes explain the emergence of IBS in males following early life stress. Our findings show pronounced sexual dimorphism in glial gene expression and functions related to communication, inflammation, and disease susceptibility. Remarkably, maternal separation has sex-specific effects on enteric glia that cause a phenotypic switch in male glia toward a phenotype normally observed in females. This phenotypic transformation is followed by physiological changes in the gut, mirroring those observed in DGBI in humans. Therefore, enteric glia appear to be a key cellular integration site in which risk factors for DGBI produce changes in gut physiology.

## RESULTS

### Early life stress induces IBS symptoms in a sex-dependent manner

Stressful events during critical periods in development, referred to here as early life stress, increase susceptibility to developing gastrointestinal disorders such as IBS, which are typified by changes in gut motor function and pain. To study mechanisms underlying these effects, we used a paradigm of maternal neglect referred to as neonatal maternal separation (NMS) stress in mice that consists of isolating pups from mothers for 3 hours per day followed by early weaning at postnatal day 17 (**Fig. 1A**). Stressed animals were compared to controls that experienced uninterrupted maternal care and a normal weaning date at postnatal day 21.

**Fig. 1.**
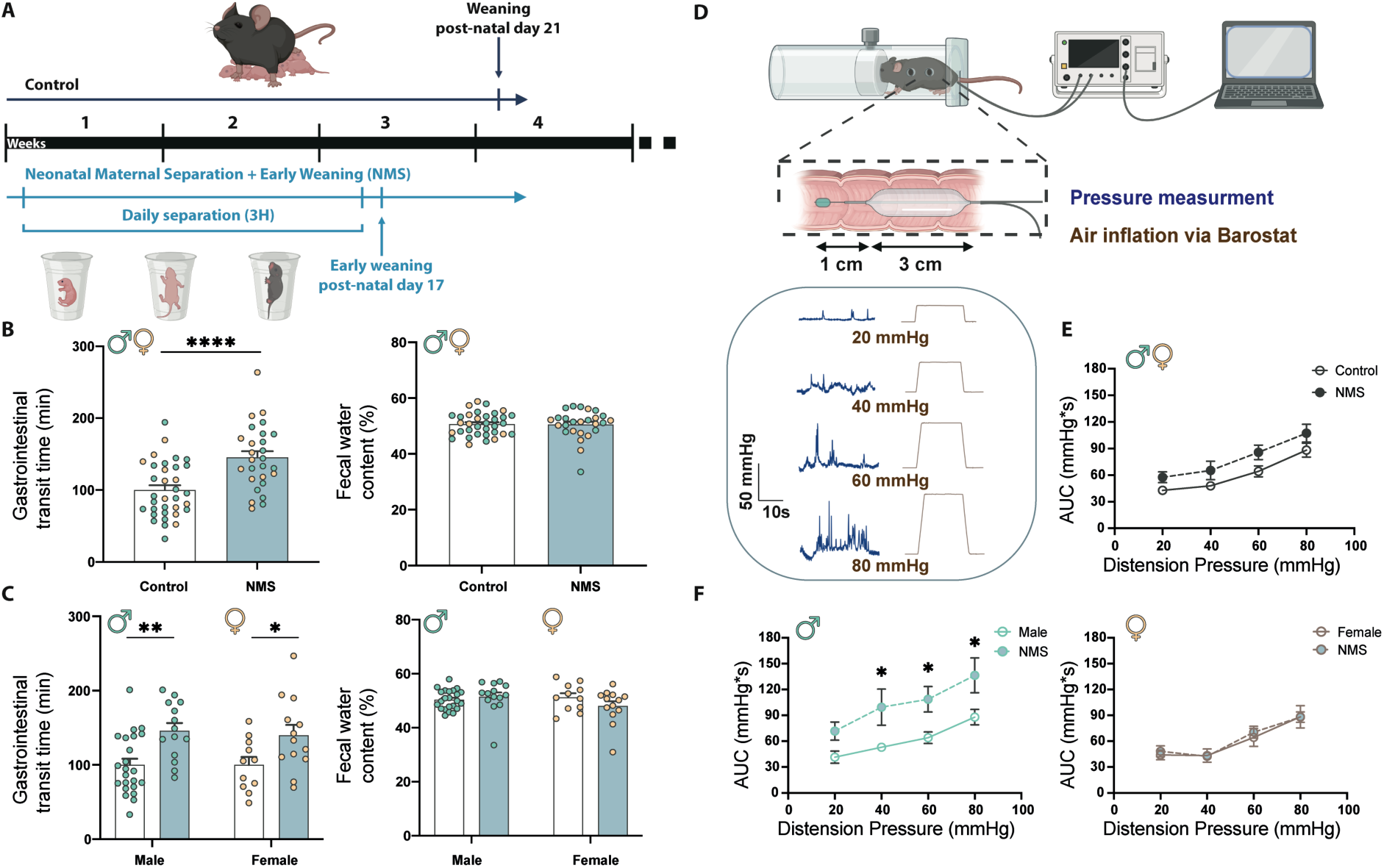
Early life stress produces IBS-like symptoms in a sex-dependent manner. **(A)** Schematic representation of the neonatal maternal separation + early weaning (NMS) paradigm. (**B**) Gastrointestinal transit time and fecal water content in 16-week controls (white) and NMS (blue) from both sexes (n=27-34), (**C**) and separated by males (green) or females (yellow) (n=11-23). (**D**) Schematic representation of visceral motor response (VMR) experiment, with representative trace for each individual pressure. (**E-F**) Visceral motor response (VMR) responses to colonic rectal distension (CRD) in mice, represented as the area under the curve (AUC) for each distension pressure. Data show VMR responses in (**E**) both sexes, (**F**) and separated by males (Left, green) and female (Right, yellow) mice in control and following NMS. Significant differences were determined by student t-test, Mann-Whitney or ANOVA 2-way followed by Bonferroni’s post-hoc analysis (* p<0.05, ** p<0.01, **** p<0.0001).

Mice that underwent NMS exhibited abnormal gastrointestinal motility that was reflected by delayed gastrointestinal transit time (**Fig. 1B**) in both males and females (**Fig. 1C** and **Table S1**). Visceral hypersensitivity is a defining feature of IBS that contributes to abdominal pain. We assessed the effects of NMS on visceral sensitivity by measuring visceral motor response (VMR) to colorectal distension (**Fig. 1D**). In agreement with earlier work^25^, control males and females have similar responses to VMR. NMS had no effect on visceral sensitivity when assessed in a mixed-sex cohort (**Fig. 1E**); however, stratifying the data by sex revealed a pronounced effect in males in which NMS increased VMR response to colorectal distension and lowered pain thresholds (**Fig. 1F**). These results show that NMS in mice recapitulates key aspects of IBS disease susceptibility in humans in which maternal neglect promotes gastrointestinal dysfunction in a sex-dependent manner^9^.

### Enteric glia are sexually dimorphic cells

We reasoned that enteric glia are an excellent candidate to mediate the effects of NMS on gut motility and sensations, given recent data showing that enteric glia modulate gut motor neurocircuits^14,15^, visceral afferent nerve terminals^16^, and are targets of psychological stress^17^. However, whether enteric glial mechanisms exhibit sexual dimorphisms that could explain the differential susceptibility of males and females is unknown. We began exploring the potential contributions of enteric glia by conducting an RNA-sequencing analysis of actively translating glial mRNA (**Fig. 2A**). Initial characterization of samples from males and females revealed stark contrasts in enteric glial gene profiles between sexes (**Fig. 2B**). Genetic glial sexual dimorphism was mainly driven by genes that were enriched in female glia with relatively few genes repressed compared to male glia (**Fig. 2C**). Genes linked to sex chromosomes were not the major driver of glial sexual dimorphism and male and female gene profiles remained distinct when sex-linked genes were excluded (data not shown). Genes differentially expressed in female glia were related to gastrointestinal, neurological, and inflammatory diseases, and represented an enrichment of several canonical pathways, including G-protein coupled receptor (GPCR) signaling, calcium signaling, endocannabinoid synaptic pathway, and S100 family signaling (**Fig. 2D**). These data agree with prior physiological data showing that female enteric glia exhibit larger magnitude Ca^2+^ responses than those in males^15^. Therefore, enteric glia are sexually dimorphic cells and female glia appear poised to contribute to inflammatory disease processes through intercellular signaling mechanisms.

**Fig. 2.**
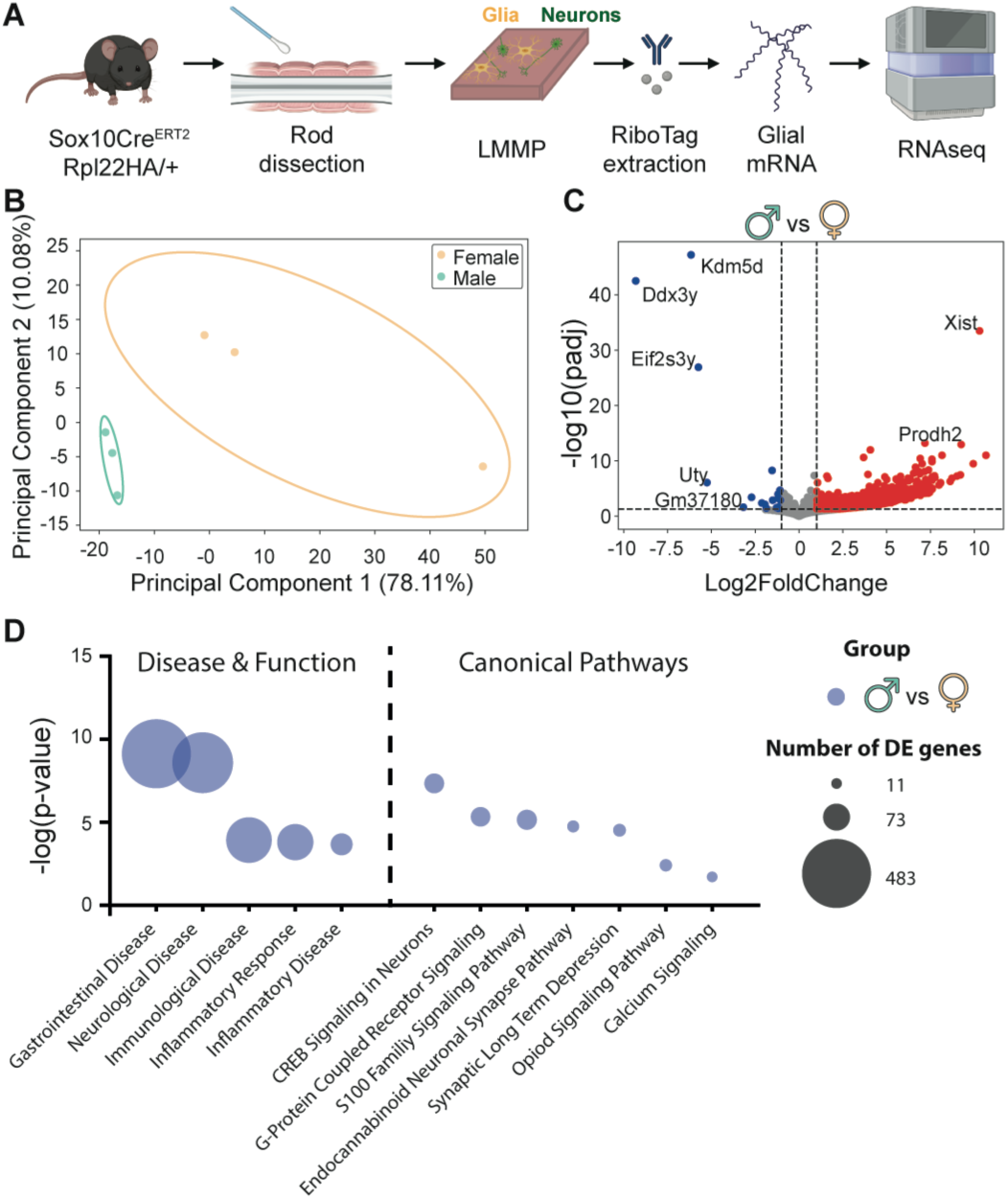
Enteric glia are sexually dimorphic. (**A**) Schematic representation of experimental design for RNAseq analysis of enteric glia. (**B**) Principal components analysis (PCA) of variance stabilized transformed counts for male and female samples (n=3 per group). Bounding ellipses were drawn based on the calculation of eigenvalues for a covariance matrix. **(C)** Volcano plot of differential gene expression between male and female enteric glia cells. Genes were considered differentially expressed when |log2fold-change| ≥ 1 and adjusted p-value ≤ 0.05. Dashed lines represent the significance threshold for fold-change and adjusted p-value. (**D**) Comparison of gene expression in males compared to females in significant pathways from Ingenuity Pathway Analsysis (IPA) ‘Diseases and functions” including GI, neurological and inflammatory disease and ‘Canonical Pathways’ including ENS signaling. The left axis represents the -log(p-value), while the size of the dot represents the number of differentially expressed (DE) genes. Genes used in IPA analysis were restricted to |log2fold-change| > 1 and FDR <0.05 (n=3 per group).

### Early life stressors induce a sexually dimorphic switch in glial genes

We repeated the transcriptomic characterization of enteric glia in males and females following NMS to explore how glial sexual dimorphism might contribute to the effects of early life stress. To our surprise, NMS had little effect on female glial gene profiles but induced a drastic shift in male glial gene expression patterns (**Fig. 3A,B**) that produced a gene expression profile similar to that of non-stressed females (**Fig. 3A-C**). In total, 786 genes were differentially regulated in male glia following NMS while only 40 were altered in females (**Fig. 3D**). These genes were distinct and only one gene (*Rnu11*) was differentially expressed following NMS in both sexes. However, genes upregulated in male glia following NMS overlapped with genes exhibiting higher expression in non-stressed females (**Fig. S1A**) and were related to the same diseases, functions, and canonical pathways as those that defined sexual dimorphism in normal controls (**Fig. 3D**). Interestingly, genes related to GPCR signaling were enriched in male glia following NMS while being reduced in females (**Fig. 3E,F**). In contrast, genes related to inflammatory signaling were reduced in males and unaffected in females following NMS (**Fig. S1B**).

**Fig. 3.**
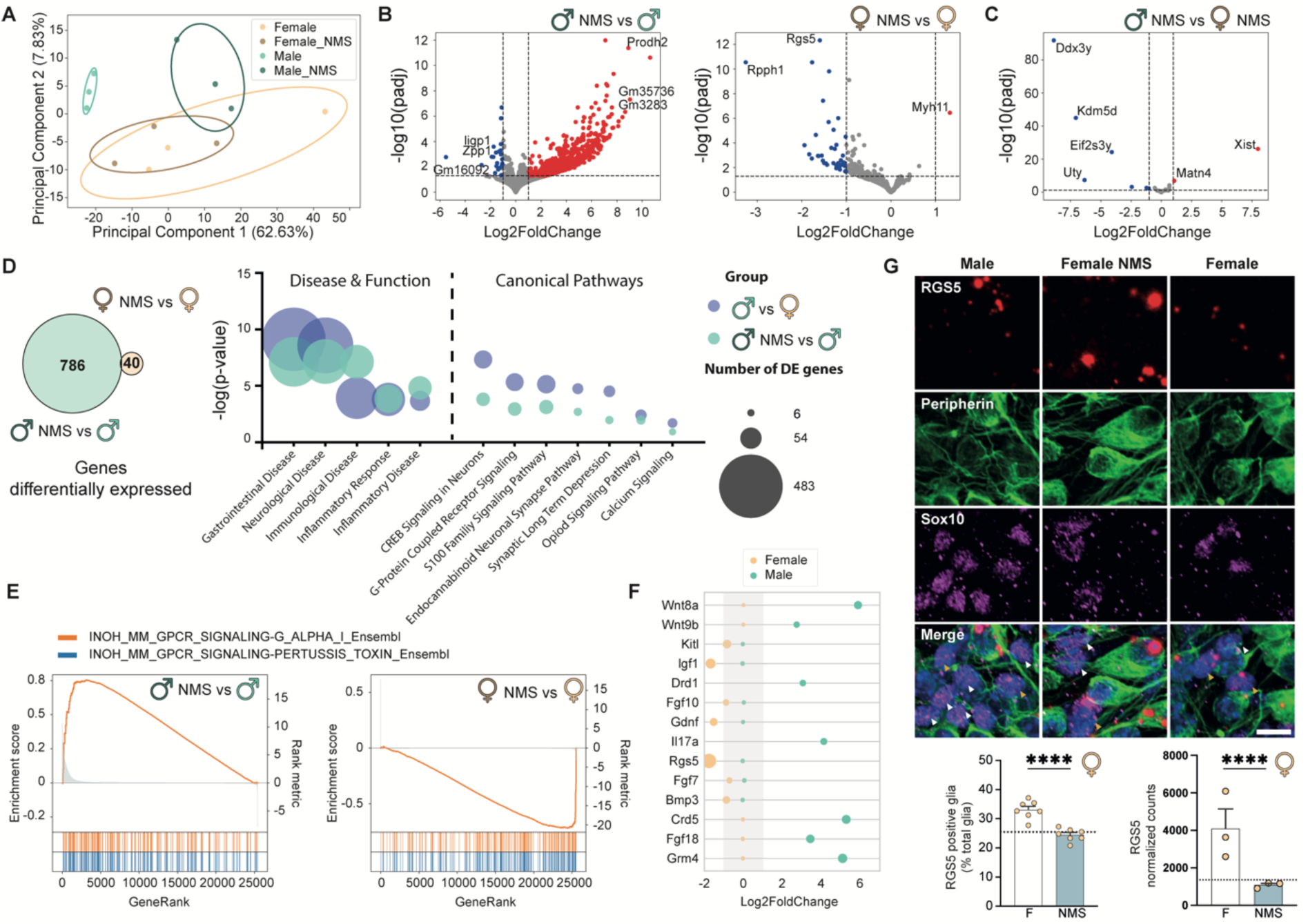
Male and female enteric glia converge on a similar genetic phenotype following early life stress. (**A**) Principal components analysis (PCA) of variance stabilized transformed counts for male and female samples from the control and NMS groups (n=3 per group). Bounding ellipses were drawn based on the calculation of eigenvalues for a covariance matrix. (**B-C**) Volcano plot of differential gene expression in enteric glia between males following NMS and control males ((**B**) left), females following NMS and control females ((**B**) right), and males following NMS and females following NMS (**C**). Genes were considered differentially expressed when |log2fold-change| ≥ 1 and adjusted p-value ≤ 0.05. Dashed lines represent the significance threshold for fold-change and adjusted p-value. (**D**) Number of genes differentially expressed in males following NMS (green) and females following NMS (yellow). Comparison of gene expression in control males compared to control females (blue dot) and males following NMS compared to control males (green dot) in significant pathways from IPA ‘Diseases and functions” including GI, neurological and inflammatory disease and ‘Canonical Pathways’ including ENS signaling. The left axis represents the -log(p-value), while the size of the dot represents the number of differentially expressed (DE) genes. Genes used in IPA analysis were restricted to |log2fold-change| > 1 and FDR <0.05 (n=3 per group). (**E**) Gene set enrichment analysis of top GPCR signaling gene sets and (**F**) gene-wise differential expression from this gene set in males (green) and females (yellow) following NMS. (**G**) RNAscope representative images of RGS5 mRNA expression in control males, females, and females following NMS. The white arrows point to non-positive glial cells, while the orange arrows point to positive glial cells for RGS5 (scale = 10µm). (Left) Quantification of positive enteric glia for RNAscope labeling and (Right) RNA seq results of *Rgs5* normalized-counts in control (white) and NMS (blue) females. The dashed line represents the expression level of control males and RNAscope quantification represents an average of at least 500 enteric glial cells per animal (n= 6-7 animals per group).

Select genes identified in the RNAseq analysis were validated using RNAscope in situ hybridization combined with immunolabeling in whole-mount samples of myenteric plexus. Given the well-established link between inflammation and DGBI pathophysiology, we focused on transcripts associated with interferon signaling and RGS5, a molecule related to GPCR signaling that regulates proinflammatory responses in astrocytes^26^. In agreement with RNAseq data, *Rgs5* expression was reduced in female glia following NMS to levels that corresponded with those observed in normal males, despite an increase in surrounding neurons (**Fig. 3G**). Similarly, interferon signaling genes (*Oasl2*, *Ifit1*, *Ifit3,* and *Gbp3*) were increased in male glia following NMS to levels that aligned with expression in control female enteric glia and were unaltered in female glia following NMS (**Fig. S1B,C**). Together, these data show that experiencing NMS causes male enteric glia to adopt a genetic phenotype similar to that of normal female glia that is poised to contribute to gastrointestinal disease.

### Early life stressors produce sexually dimorphic alterations in glial signaling

Sex- and stress-related differences of glial genes related to cellular communication and GPCR signaling pathways suggest that changes to glial signaling mechanisms, which are central modulators of ENS activity and gut functions^14,16,27^, could contribute to abnormal motility and pain following NMS. We studied changes to glial activity using a dual transgenic system in which enteric neurons and glia express the genetically encoded calcium indicator GCaMP5g, and enteric glial activity is controlled via the chemogenetic receptor hM3Dq^14,15^. Focal stimulation of ganglia with the hM3Dq agonist clozapine-N-oxide (CNO) in whole-mount preparations of the colon myenteric plexus provided targeted activation of glia within specific ganglia, and subsequent effects on glial and neuronal activity were observed using GCaMP imaging (**Fig. 4A**). In agreement with earlier work^15^, glial response to CNO were sexually dimorphic and female enteric glia exhibited larger calcium responses, a shorter peak duration, and lower frequency of response compared to male enteric glia (**Fig. 4B-F**). Similar to the effects on gene expression profiles, recordings of cells from animals that underwent NMS showed that this normal difference in glial responsiveness between males and females was reduced with glia converging on a similar phenotype. NMS decreased glial Ca^2+^ response intensity (24.6% decrease in males and 23.1% in females) (**Fig. 4C**) and increased response frequency post-stimulation (27.6% increase in males and 77.2% in females) in both sexes (**Fig. 4E,F**) while showing opposite-sex effects on response duration which were shorter in males (9.9% shorter) and longer in females (9.6% longer) (**Fig. 4D**). These results show that NMS causes male and female glia to adopt a similar physiological phenotype through changes in patterns of cellular activity and responsiveness.

**Fig. 4.**
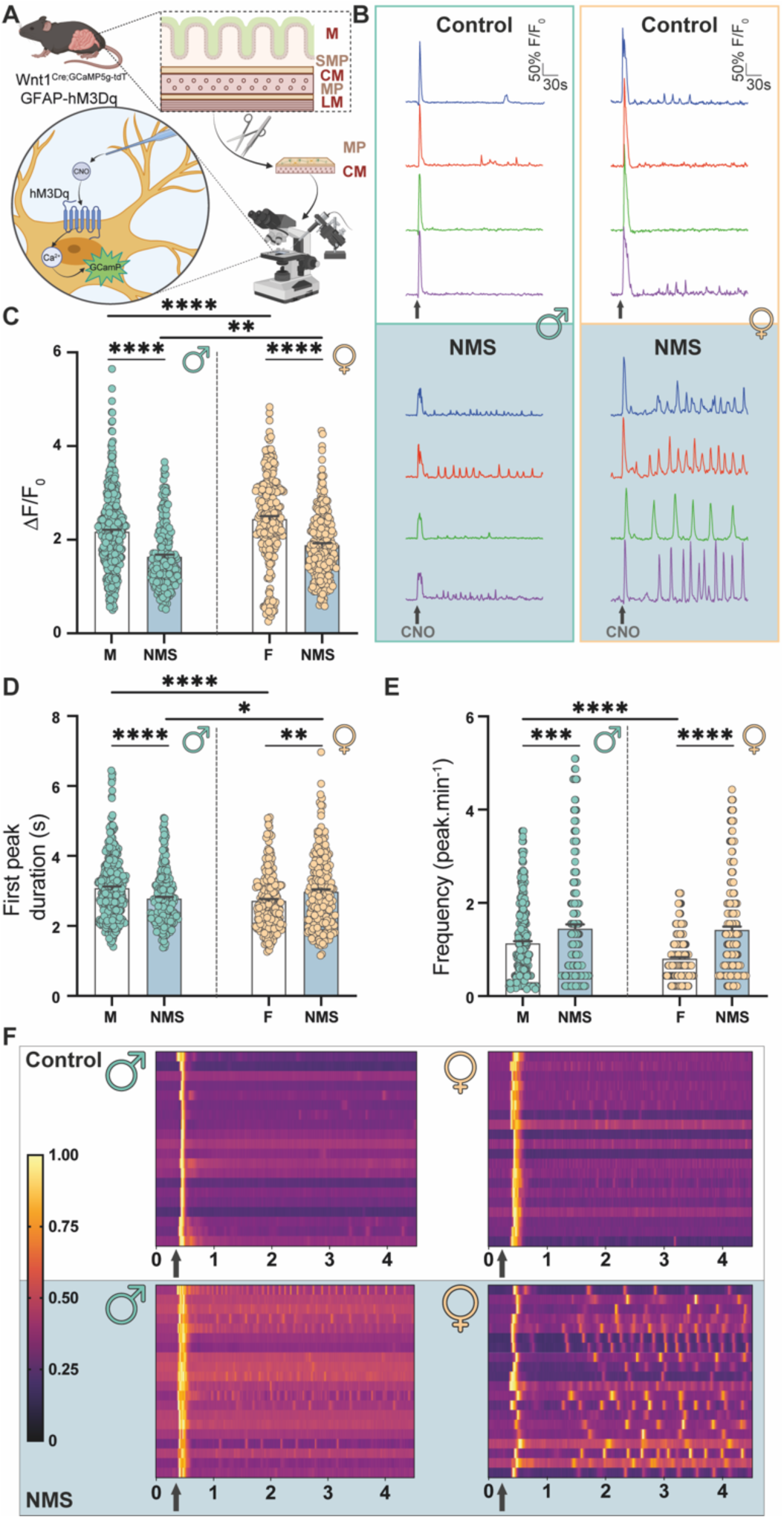
Early life stress disrupts physiologic enteric glial activity. (**A**) Schematic methodology of the experiment (M: Mucosa; SMP: Submucosal plexus; CM: Circular muscle; MP: Myenteric plexus; LM: Longitudinal muscle). (**B**) Representative traces reported as ΔF/F_0_ over time of individual cell’s calcium response following a CNO puffing stimulation (arrow) in control males (Top left) and females (Top right), NMS males (Bottom left) and females (Bottom right). (**C-E**) Quantification of maximal peak ΔF/F_0_ response (**C**), first peak response duration and frequency of peaks (**E**), following CNO stimulation in control (white) and NMS (blue) males (green dots) and females (yellow dots) animals. (**F**) Heat map of 20 enteric glia ΔF/F_0_ response over time; from a representative ganglion following CNO stimulation (arrow) in control (top, white) and NMS (bottom, blue) males (left) and females (right). Graph bars represent mean + SEM, and each individual dot represents the value of one glial cell. Values for each group were analyzed via two-way ANOVA followed by Bonferroni’s post hoc analysis (* p< 0.05, ** p< 0.01, *** p< 0.001, **** p< 0.0001, n = 234-390 cells from 3-5 animals per group (from at least 3 different ganglia per animal)).

The changes to glial activity observed following NMS could be due to direct effects on glia or indirect effects mediated by changes to enteric neurons^28–30^ that communicate with enteric glia^27^. To assess potential neuronal contributions to changes in glial Ca^2+^ responses driven by CNO, we incubated samples with TTX to block neuronal signaling downstream of action potentials (**Fig. S2A**). TTX did not affect glial peak response amplitude to CNO in control animals but decreased responses in males and increased responses in females following NMS. Glia also exhibited increased response frequency when neuronal activity was reduced in non-stressed males and females (**Fig. S2B-D**), suggesting the removal of an inhibitory neuronal input to glia. This effect was most pronounced in males, in which we observed an increase in the frequency of glial Ca^2+^ responses and shorter duration glial responses following NMS and neuronal action potential blockade (**Fig. S2B,C**). Neuronal inhibition of glial response intensity in females was more pronounced post-NMS (**Fig. S2B,D**). However, blocking neuronal input did not entirely suppress the difference observed between non-stressed and stressed animals, indicating that NMS alters neuron-glial communication through effects on both neurons and glia.

### Changes to glial signaling contribute to sex-specific changes in gut function following early life stress

Enteric glial activity encoded by Ca^2+^ responses modulates gut neurocircuits that control intestinal motility^14^. Given that experiencing NMS disrupts this mechanism of glial activity, we speculated that changes to glial signaling contribute to abnormal intestinal motility following NMS. We began testing potential differential effects of glial activity on gut motility by measuring gastrointestinal transit time (GITT) following glial stimulation with CNO in hM3Dq^+^ mice. GITT was delayed in both sexes following NMS, with no discernible impact on fecal water composition (**Fig. 5A-F** and **Table S1**). Stimulating glial activity did not affect total GITT in control animals^14^, but normalized transit times toward controls in NMS mice (**Fig. 5A**). Splitting the data by sex revealed that this effect was driven by stressed males which showed a return to normal GITT following glial activation (**Fig. 5B**). In contrast, glial stimulation did not affect the delay in GITT observed in NMS females (**Fig. 5C**). These data show that the switch in male glial phenotype following NMS produces physiologically relevant effects on gut function in males and that the minor effects on enteric glia in females do not make major contributions to altered gut motility in NMS females.

**Fig. 5.**
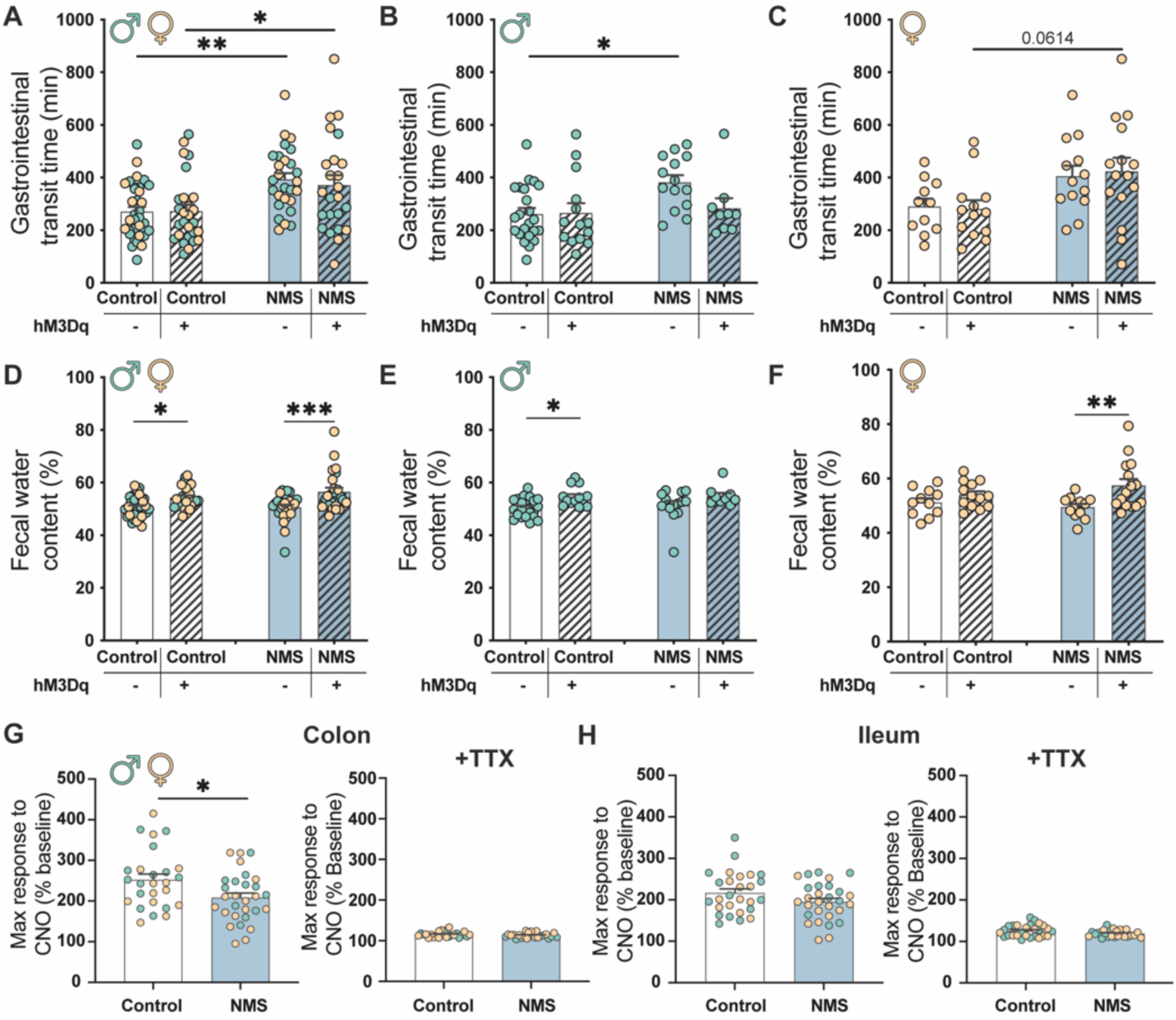
Early life stress alters gastrointestinal motility through a change in enteric glia. (**A-C**) Gastrointestinal transit time in control or NMS animals without or with the expression of the chemogenetic receptor hM3Dq in enteric glia (hM3Dq(–) and hM3Dq^(+)^ respectively) from (**A**) both sexes, (**B**) males only and (**C**) females only (n= 9-23). (**D-F**) Fecal water content from (**D**) both sexes, (**E**) males only and (**F**) females only. (**G-H**) Organ isolated bath experiment showing the contractile response to CNO from colon (**E**) and ileum (**F**) without or with the presence of tetrodotoxin (**G-H**) (n= 25-29). Graph bars represent mean + SEM, and each individual dot represents the value of one animal. Significant differences were determined by Student t-test or Two-way ANOVA followed by Bonferroni’s post-hoc analysis (* p<0.05, ** p<0.01, *** p<0.001).

Stimulating glia with CNO in organ bath experiments using samples of colon and ileum from hM3Dq^+^ mice recapitulated prior results showing that glial activity promotes gut contractions in both males and females^14,31^ (**Fig. 5G,H**). Interestingly, the pro-contractile effect of glia was reduced in the distal colon of NMS mice of both sexes (**Fig. 5G** and **Table S3**) but was not altered by NMS in the ileum (**Fig. 5H**). Blocking neuronal action potentials with TTX eliminated contractions produced by electric field stimulation (**Table S2**). TTX reduced colonic and ileal contractions following CNO stimulation and removed the difference between controls and NMS mice (**Fig. 5G,H**). These results indicate that changes in motility observed after NMS involve indirect communication between enteric glia and muscle cells through neuronal pathways. These results suggest that changes produced by glial stimulation are region-specific and agree with results presented earlier showing pronounced effects in colonic glia.

### Changes to prostaglandin E2 signaling contribute to abnormal glial activity and motility in males following NMS

The data indicate that experiencing NMS induces robust changes in enteric glial phenotype that contribute to abnormal motility in males and pathway analysis suggests that the male NMS glial phenotype may involve an upregulation of proinflammatory properties. We recently showed that enteric glia upregulate prostaglandin E2 (PGE2) during inflammation and that glial PGE2 sensitizes visceral sensory neurons in females^16^. Based on these results, we hypothesized that changes in PGE2 signaling might contribute to abnormal glial activity and motor responses following NMS in males. Given that glial Ca^2+^ activity decreased following NMS, we initially focused on effects of PGE2 mediated through EP3 receptors (**Fig. 6A**). EP3 receptors predominantly couple to inhibitory G_i_ signal transduction pathways; however, in related glia such as astrocytes, EP3 receptors couple to pathways that enhance Ca^2+^ responses^32^. Similarly, we found that blocking EP3 receptors with L-798,106 reduced enteric glial Ca^2+^ response magnitude in control males and females to a similar extent (**Fig. 6B,C**). However, the effect of blocking EP3 receptors was amplified in male glia following NMS while no additional effect was observed in females (**Fig. 6B,C**). L-798,106 also produced a robust increase in Ca^2+^ response frequency in male glia following NMS but had no effect on glial response frequency in normal or NMS females.

**Fig. 6.**
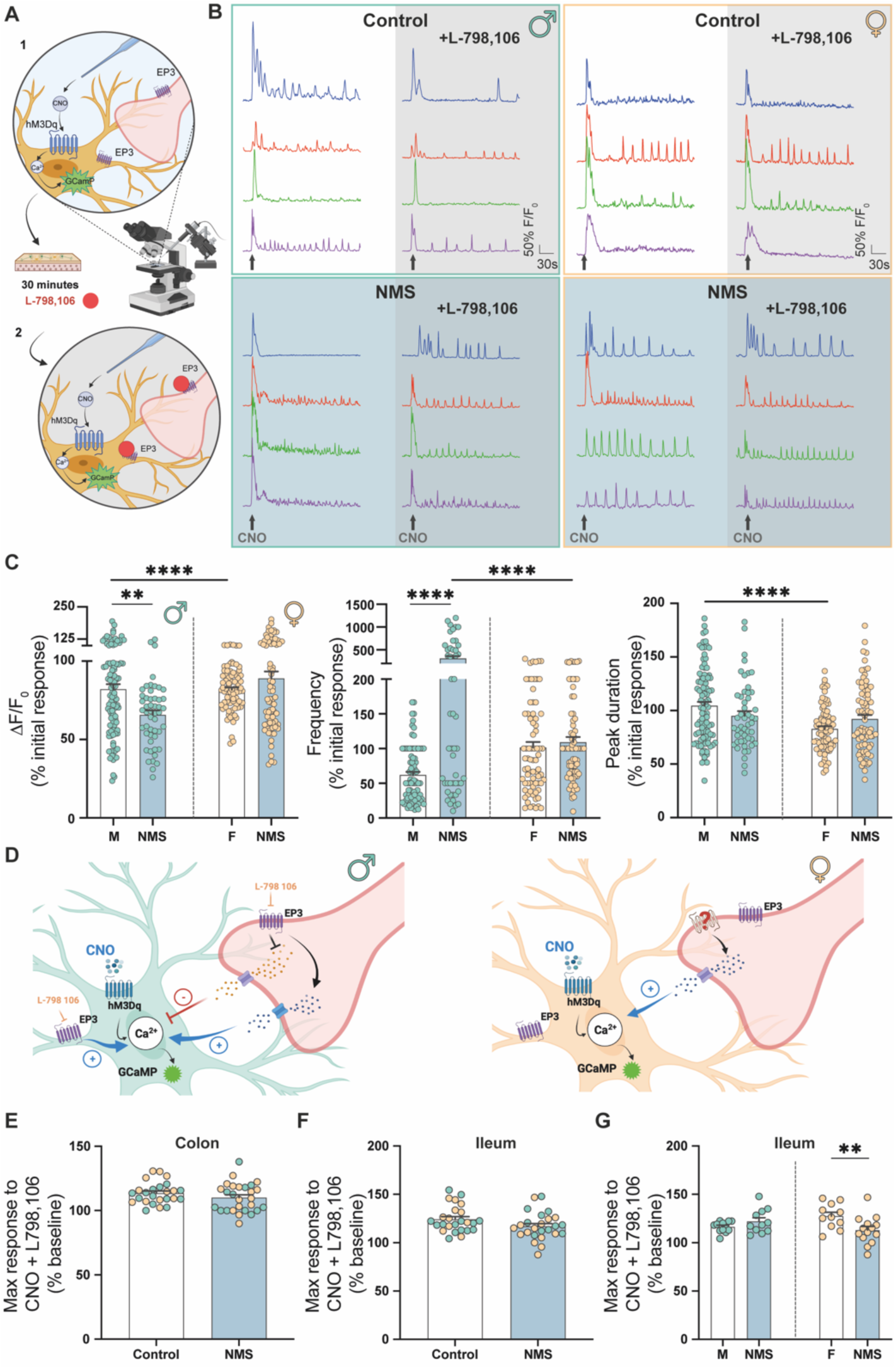
Prostaglandin E2 signaling contributes to abnormal glial signaling and motility following early life stress in males. (**A**) Schematic methodology of the experiment. (**B**) Representative traces reported as ΔF/F_0_ over time of individual cell’s calcium response following a CNO puffing stimulation (arrow) in control males (Top left) and females (Top right), NMS males (Bottom left) and females (Bottom right) in the absence (white) or in the presence of L-798,106 (grey), a specific PGE2 receptor 3 antagonist. For each condition, color-matched traces represent a glial cell response to CNO before and after L-798,106 treatment. (**C**) Quantification of maximal peak ΔF/F_0_ response, frequency of response, and maximal peak response duration following CNO stimulation in control (white) and NMS (blue) males (green dots) and females (yellow dots) animals (n = 52-100 cells from 2-4 animals per group (from 1 ganglion per animal)). (**D**) Schematic representation of proposed mechanisms altered after NMS in male (top) and female (bottom) enteric glia. (**E**) Colonic and (**F**) ileal contractile response to CNO stimulation in the presence of (top) L-161,982, an EP4 antagonist, and (bottom) L-798,106, an EP3 receptor antagonist. (**F** right) Response to CNO response in the presence of EP4 or EP3 receptor antagonists separated by biological sex. Graph bars represent mean + SEM, and each individual dot represents the value of one glial cell. Values for each group were analyzed via two-way ANOVA followed by Bonferroni’s post hoc analysis (* p< 0.05, ** p< 0.01, *** p< 0.001, **** p< 0.0001).

To investigate how changes in glial activity mediated by PGE2 contribute to motor dysfunction following NMS, we repeated organ bath experiments in the presence of L-798,106. Prior data indicated that EP4 receptors also modulate motor responses in the mouse colon and ileum^33^, so we also conducted studies with the EP4 antagonist L-161,982 (**Table S4**). Blocking either EP3 or EP4 receptors eliminated the excitatory effect of glial stimulation in both the ileum and colon and removed the decrease in motor function following NMS (**Fig. 6E,F**). Splitting the data by sex revealed that tissue from NMS females still exhibited a decreased motor response to glial stimulation in the presence of either EP3 or EP4 antagonists but the effect in males was lost (**Fig. 6G and Table S4**). Therefore, changes to PGE2 signaling in males contribute to the effects of NMS on glial activity and motor responses.

### Glial mechanisms underlying heightened visceral sensitivity in males following NMS

Visceral hypersensitivity is a major mechanism underlying pain in IBS. Our data show that experiencing NMS promotes visceral hypersensitivity in males (**Fig. 1**) and that this effect is accompanied by a phenotypic switch in male glia that involves abnormal signaling through pain-related mechanisms such as endocannabinoid and opioid pathways. To investigate glial mechanisms involved in this switch that sensitize dorsal root ganglion (DRG) sensory neurons, we performed an interactome analysis between ligand genes enriched in enteric glia following NMS with receptor genes expressed by colon-projecting DRG neurons^34^. Thirty-seven glial ligand-encoding genes emerged as candidates that were paired with 32 corresponding receptors in colon-innervating DRG sensory neurons (**Fig. 7A-C** and **S3A-D**). DRG neuron subtypes were further segregated into nociceptive (mNP, mPEPa, mPEPb and pPEP) and non-nociceptive populations (mNFa, mNFb and pNF) (**Fig. 7B** and **S3B**). Strikingly, there was little similarity between differentially expressed glial ligands following NMS in males and females (**Fig. 7C** and **S3C**). As an example, interactions between enteric glia and mNP nociceptors mediated by hyaluronan derived from glial hyaluronan synthase (HAS2) and nociceptor CD44 receptors are predicted to be enhanced following NMS in males (**Fig. 7A** and **S3A**). CD44 is involved with neuronal and glial guidance and synaptic transmission^35^ and abnormal CD44 signaling could contribute to the observed effects on visceral hypersensitivity NMS males. Other upregulated genes in male enteric glia included *PNOC* and *CYP19A1*. *PNOC* encodes nociception, a neuropeptide with pain-modifying effects^36,37^, while *CYP19A1* encodes aromatase, an enzyme that converts androgens to estrogens. Its product, estradiol, regulates inflammation and increases visceral sensitivity by altering TRPV1 expression^38,39^. These data support the concept that NMS alters specific mechanisms of interaction between glia and nociceptive neurons in males that likely contribute to increased visceral sensitivity following NMS.

**Fig. 7.**
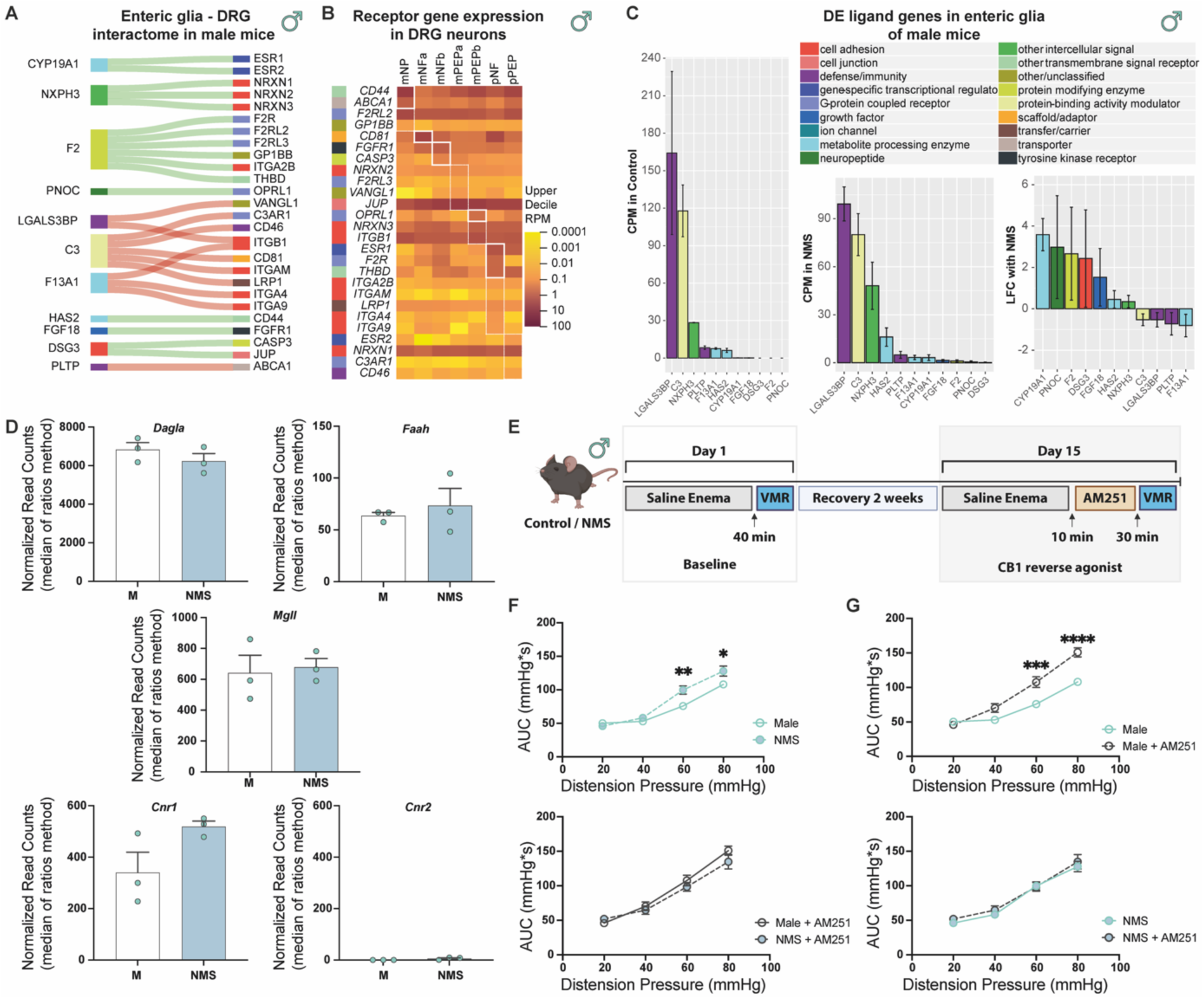
Gene expression patterns predict mechanisms of interaction between enteric glia and dorsal root ganglion neurons that contribute to visceral hypersensitivity following early life stress in males. (**A**) Ligand genes in male enteric glia that were differentially expressed with NMS (adj. p-value <0.05) and their receptor pairs that were expressed in DRG neurons. Green flows indicate upregulation of the ligand gene, and red flows indicate downregulation following NMS. (**B**) Heatmap of receptor gene expression in different populations of colon-innervating DRG sensory neurons; the thick white box indicates enrichment of the gene in the corresponding neuronal population, and the thin dotted box indicates the population with the highest expression of a nonspecific gene. (**C**) Expression of the ligand genes in control and NMS enteric glia (error bars indicate standard deviation) and log2 fold-change with NMS (error bars indicate standard error). NMS = neonatal maternal separation (n=3); Control (n=3). (**D**) RNA seq results of endocannabinoid signaling-related genes *Dagla, Faah, Mgll, Cnr1 and Cnr2* normalized-counts in control (white) and NMS (blue) males (n=3). (**E**) Experimental timeline for Visceral motor response at baseline and with cannabinoid receptor 1 (CB1) reverse agonist AM251. (**F**) Visceral Motor response (VMR) responses to Colonic Rectal distension (CRD) in mice, represented as the area under the curve (AUC) for each distension pressure. Data show (Top) baseline VMR responses and (Bottom) then with injection of cannabinoid receptor 1 reverse agonist AM251 in control and NMS animals (n=11-12). (**G**) VMR response in (Top) control and (Bottom) NMS animals with or without AM251 (n=11-12). Significant differences were determined by ANOVA 2-way followed by Bonferroni’s post-hoc analysis (* p<0.05, ** p<0.01, **** p<0.0001).

As a proof of concept, we chose to focus on potential contributions of endocannabinoid-mediated glial-DRG interactions in visceral hypersensitivity following NMS based on observed alterations to glial endocannabinoid pathways in Ingenuity Pathway Analysis (IPA) analysis between sexes and after NMS in males (**Fig. 3D**) and known roles of endocannabinoids in modulating intestinal motility and visceral sensitivity^40^. While changes in enzymes responsible for producing or degrading endocannabinoids did not reach significance in either sex, glial cannabinoid receptor 1 (CB1, encoded by *Cnr1*) mRNA expression increased in males following NMS (**Fig. 7D and S3C**), which agree with clinical observations suggesting *Cnr1* polymorphisms at the locus rs806378 and variation in the number of AAT triplet repeats are associated with changes in CB1 expression in IBS patients^41^. To understand the potential contributions of altered glial CB1 receptor expression to male visceral hypersensitivity following NMS, we assessed how blocking CB1 receptors with the inverse agonist AM251^42^ affects NMS-driven visceral sensitivity in males (**Fig. 7E**). Interestingly, blocking CB1 with AM251 increased visceral sensitivity in control males to levels observed following NMS (**Fig. 7F,G**), which would support the idea that normal endocannabinoid signaling dampens visceral sensitivity. However, effects of AM251 on visceral sensitivity were lost following NMS, suggesting a disruption of baseline endocannabinoid control. Taken together with the gene expression data, these results support the conclusion that changes to endocannabinoid signaling between enteric glia and nociceptors contribute to increased visceral sensitivity in males following NMS.

## DISCUSSION

IBS and related DGBIs are common disorders with an unexplained etiology that involves sex, inflammation, and stress. Females are typically more susceptible to developing IBS, but certain early life stressors program a female-like susceptibility in males. Here, we discover that enteric glia are a key cellular integration site for IBS susceptibility cues. Enteric glia are sexually dimorphic cells that tune intestinal neurocircuits controlling motor function and pain. Importantly, experiencing early life stress induces a sex-dependent switch in glial phenotype that promotes abnormal gut motility and visceral hypersensitivity in males. Targeting these context- and sex-dependent glial mechanisms could ultimately benefit the treatment of disorders such as IBS.

Enteric glia maintain homeostasis within the ENS and modulate gut neurocircuits that control motility, epithelial functions, and visceral sensitivity^43^. These actions appear to result from discrete subsets of enteric glia that have specialized functions within neural networks and in spatially diverse compartments of the gut wall^15,44,45^. Despite an appreciation that extensive glial diversity exists in the gut, the extent to which glial functions are dictated by biological sex has remained unknown. Earlier work provided hints that enteric glia exhibit sexually dimorphic functions by showing differential effects of glial ablation on gut motor function in females^46^, sex differences in glial activity encoded by Ca^2+^ responses^15^, and differing roles in visceral pain induced by inflammation^16^. Our data show that sex-specific glial gene profiles contribute to these functional differences. Similar genetic sex differences are thought to contribute to differential roles of human and rodent astrocytes and microglia^47–49^. The differential gene expression across sexes is linked to baseline differences in glial function, which alters their Ca^2+^ response to external stimuli and modulates their response to inflammation or injury^49,50^. Glial sexual differences are also altered by lifestyle elements, such as psychological stressors, changes in diet or microbiome which ultimately change glial functions in the brain^51^. Similarly to this observation, our data demonstrate that the gene differences in enteric glia are associated with differences in glial cellular activity and that the response to external stressors in enteric glia is unique for each biological sex.

Multiple stressors contribute to early life stress such as neglect, poverty, and family bereavement. The impact of early life stress on the gastrointestinal tract varies with the type of stressor^25^ with context-specific effects on various cellular populations that contribute to intestinal disorders^52^. Previous studies demonstrate that the nature of the stressor is important and can differentially affect intestinal functions in each sex^9,25,52^. Enteric glia act as transducers in the scenario of psychological stressors, which lead to changes in intestinal function^17^. Enteric glia are still in the process of maturation during the early postnatal period^53^ and repeated stress alters their morphological phenotype and functions into adulthood^17,24^. The molecular underpinnings involve gene changes that reduce sexual dimorphism in enteric glia and lead to DGBI-related intestinal dysfunction. This switch mirrors the changes observed in central glia following early life stress characterized by an increase of glial activity and a switch to reactive glia^54^. These alterations might emerge from early life stress-induced epigenetic changes that affect gene expression and gastrointestinal functions^55–57^, or changes in master transcription factors which modify several downstream genes in their signaling pathways^58^.

Experiencing early life stress increases the risk of developing DGBI in adulthood that manifest as abnormal intestinal motility and visceral hypersensitivity^2,4^. These symptoms are associated with abnormal glial interactions with enteric neurons and/or immune cells with subsequent impacts on intestinal functions^15,43,59^. Our findings demonstrate that early life stress disrupts enteric glial phenotype and that these changes impact intestinal transit. Remarkably, targeted stimulation of enteric glia corrects motility in animals subjected to early life adversity. Ablation models suggest that enteric glia influence intestinal motility mainly in females^46^, and the shift in male glia to a female-like phenotype participates in intestinal dysmotility after early life stress. The phenotype change induces an alteration of glial mediators signaling. One of these mediators is prostaglandin E2 (PGE2) which modulates intestinal and ENS function^60–63^ through its receptors (EP1 to EP3) in homeostasis and inflammation^16,33,64,65^. After early life stress, alterations of the PGE2-EP3 signaling occur at different levels in each sex. These changes affect glia-to-glia communication within the myenteric plexus in males, while it affects glia-to-neurons and muscle communication in females^66^. Considering that enteric glia upregulate PGE2 secretion under inflammatory conditions^16^ and undergo a phenotypic shift following early life stress, it is likely that the alteration in glial PGE2 secretion contributes to the observed changes in motility and glial activity.

Enteric glia modulate visceral sensitivity through mechanisms that involve communication with immune cells^18^ and direct interactions with nociceptive nerve terminals^16^. Following exposure to early life stress, males develop visceral hypersensitivity^52^. This is associated with changes in gene expression that link ligand-receptor interactions between enteric glia and dorsal root ganglia, which transduce information related to visceral sensitivity^40^. Of the multiple pathways facilitating communication between enteric glia and neurons, endocannabinoid signaling emerged as a promising mechanism that was altered in males after early life stress. Endocannabinoid signaling provides a balance of inhibitory control over signaling within the ENS and plays a major role in modulating visceral sensitivity^40^. In the CNS, the cannabinoid receptor 1 activation induces a direct inhibition of neuronal activity and an increase in astrocytic Ca^2+^ levels, thereby enhancing glial communication^67^. In the ENS, changes in glial endocannabinoid signaling were evident following early life stress and indicated potential disruptions in the control of glia and neuronal pathways. In support, blocking cannabinoid receptor 1 increases visceral hypersensitivity in control males. Interestingly, blocking cannabinoid receptor 1 following early life stress does not alter the VMR response, despite a noticeable increase in cannabinoid receptor 1 mRNA expression in males. This suggests that increased visceral sensitivity following early life stress may lead to desensitization of the cannabinoid receptor 1^68^, resulting in a reduced analgesic response and a compensatory increase in mRNA expression. Alternatively, limitations of the VMR technique may contribute to this observation, as the visceral response following early life stress might reach the detection limit and lead to non-detection of an increased response. However, our results argue for a role of endocannabinoid signaling in abnormal enteric glial-neuron interactions that occur following early life stress and suggest its potential therapeutic relevance in males. Further studies are necessary to understand the complexities of endocannabinoid signaling pathways in this context and the extent to which enteric glia are involved.

The understanding of sex-specific responses to stress in the context of gastrointestinal disorders presents a challenge with significant clinical implications. Despite females traditionally exhibiting higher susceptibility to these conditions, our results demonstrate that certain early life stressors induce a female-like susceptibility in males. Here, we discover that enteric glia represent a key cellular integration site predisposing individuals to functional gastrointestinal disorders. Enteric glia regulate gut motility and pain perception, and alterations in their phenotype contribute to abnormal gastrointestinal functions. By elucidating the distinct genetic profiles of enteric glia across sexes, this study sheds light on the mechanisms underlying their functional diversity. Moreover, our findings offer profound insight into the impact of early life stress on glial functions, ultimately leading to an IBS-like phenotype. This study paves the way for novel therapeutic strategies targeting sex-specific mechanisms underlying DGBI, thereby offering improved management and treatment for these conditions.

## MATERIALS AND METHODS

### Animals

All work involving animals was conducted in accordance with the National Institutes of Health (NIH) *Guide for Care and Use of Laboratory Animals* and was approved by the Institutional Animal Care and Use Committee (IACUC) at Michigan State University (AUF# PROTO202100150) in specific pathogen-free conditions in facilities accredited by the Association for Assessment and Accreditation for Laboratory Care (AAALAC) International. An equal number of male and female mice with *C57BL/6J* backgrounds were used for experiments unless otherwise stated. Animal age is stated in each subsection of the materials and methods. All animals were maintained on a 12h:12h light-dark cycle in a temperature-controlled environment (Optimice^®^ cage system; Animal Care Systems, Centennial, CO) with access to minimal phytoestrogen diet (Diet Number 2919; Envigo, Indianapolis, IN) and water *ad libitum*.

So×10^CreERT2+^;Rpl22^tm1.1Psam^/J mice (referred as GliaRiboTag) were generated as described previously^69^ to express the hemagglutinin (HA) tag on ribosomes within enteric glia upon tamoxifen administration. This line was maintained as hemizygous for both Cre (MGI:5910373; So×10^CreERT2+/-^ a gift from Dr. Vassilis Pachnis^70^, The Francis Crick Institute, London, England) and the floxed allele (Rpl22^tm1.1Psam^/J; Jackson Laboratory, Bay Harbor, ME, stock number 011029; RRID: IMSR_JAX:011029). Transgenic mice expressing the genetically encoded floxed allele of calcium indicator GCaMP5g (Jackson Laboratory, Bay Harbor, ME; RRID: IMSR_JAX:024477) were crossed with *GFAP-hM3Dq* (MGI:6148045; gifted by Dr. Ken McCarthy^71^; University of North Carolina, Chapel Hill, NC) and were bred for experiments to produce *GCaMP5gtdT^f/f^::GFAP-hM3Dq^+^*. These mice were crossed with *Wnt1^CreER2^* (Jackson Laboratory, stock number 022137; RRID: IMSR_JAX:022137) and were bred for experiments as heterozygotes to produce *Wnt1^Cre;GcaMP5gtdT^::GFAP-hM3Dq* mice. Triple-transgenic mice were maintained as hemizygous for Cre (*Wnt1^CreER2^*). GliaRiboTag mice were fed tamoxifen citrate (400mg.kg^-1^ diet) for 2 weeks to induce HA expression. Genotyping was performed by the Research Technology Support Facility Genomics Core at MSU and Transnetyx (Cordova, TN).

### In vivo model of Neonatal Maternal Separation and early weaning

NMS was conducted as previously described^23^ and represented in **Fig. 1A**. Briefly, C57BL/6J pregnant nulliparous dams were isolated before giving birth. The first day pups were observed was denoted as postnatal day 0 (PD0). For control groups, pups were raised under standard protocols and undisturbed in home cages until weaning at PD21. For the NMS model, mice were subjected to 3 hours of isolation (Zeitgeber +3-6) from PD1-PD16 and weaned at PD17. The separation was done by placing each pup in individual plastic cups containing a small amount of their home cage bedding. The cups were housed in separate cages with 5 cups/cage. NMS mothers were placed in separate clean cages with food and water *ad libitum* access. After 3 hours of separation, pups and dams were returned to their home cage with standard food, water, bedding, and day/night cycling. The animals were undisturbed until the next morning of separation.

### Myenteric plexus isolation

Distal colonic myenteric plexus was dissected as previously described^69^ to maintain RNA integrity. Briefly, mice were euthanized by cervical dislocation and subsequent decapitation. The distal half of colonic tissue from male and female control and NMS GliaRiboTag mice were removed and placed in a petri dish containing ice-cold Dulbecco’s Modified Eagle Medium (DMEM/F-12) surrounded by dry ice. Luminal contents were flushed with a syringe, and a plastic rod (∼3.2 mm diameter) was inserted into the lumen. The longitudinal muscle-myenteric plexus (LMMP) was isolated by gently removing the mesenteric border and teasing away the LMMP from the underlying circular muscle using cotton swabs. For RNA-seq, LMMP samples were immediately flash-frozen in 1.5 mL tubes in liquid nitrogen and stored at −80°C until further processing. For RNAscope and immunohistochemistry, LMMPs were pinned flat in a Sylgard - coated petri dish and fixed overnight in 4% paraformaldehyde (PFA) at 4°C. Fixed tissue was washed 3 × 10 minutes with 1x phosphate-buffered saline (PBS) and stored in 1x PBS at 4°C until further processing.

### RiboTag extraction

Flash-frozen distal colon LMMPs from GliaRiboTag mice were homogenized using a CryoGrinderTM (OPS Diagnostics, Lebanon, NJ) mortar and pestle cooled using liquid nitrogen. Samples were ground into a fine powder for 20-30 s and transferred to an ice-cold 1.5 mL tube. RiboTag isolation was performed on these homogenized samples as previously described with solution adjustments for gastrointestinal tissue^72,73^. Briefly, 1 mL of supplemented homogenization buffer (50 mM Tris, pH 7.4, 100 mM KCl, 12 mM MgCl_2_, 1 % Nonidet P-40, 1 mM DTT, 10 mg sodium deoxycholate, 100 μg.mL^-1^ cyclohexamide, 20 μL protease inhibitor, 10 μL RNAse inhibitor (40 U.μL^-1^), and 3 mg.mL^-1^ heparin in RNAse-free water) were added to each sample vial and mixed vigorously by pipetting. Samples were centrifuged at 10,000 *g* for 10 minutes, and ∼800 μL of supernatant was transferred to a clean 1.5 mL vial with 5 μL HA antibody. Samples were gently rotated on an Eppendorf tube rotator (Rotoflex R2000, Argos Technologies) for 4 hours at 4°C. Samples were added to 200 μL Protein A/G beads and gently rotated overnight at 4°C. The following day sample supernatants were removed using a magnetic stand on ice, and sample beads were washed 3 × 10 minutes at 4°C with high salt buffer (50 mM Tris, pH 7.5, 300 mM KCl, 12 mM MgCl_2_, 1% Nonidet P-40, 1 mM DTT, 100 μg.mL^-1^ cycloheximide)^74^ and subsequently eluted with 350 μL RLT Plus lysis buffer with DTT.

Eluted mRNA was purified using the RNeasy Micro Kit Plus and quantified using the Quant-ITTM RiboGreen® RNA Assay Kit using the low-range standards according to the manufacturer’s protocols. RNA integrity number (RIN) was assessed using a 2100 Bioanalyzer and the Eukaryote Total RNA 6000 Pico chip (Agilent, Santa Clara, CA), and samples with RIN > 6.5 were used for sequencing.

### RNA library preparation and sequencing

Libraries were prepared by the Van Andel Genomics Core (Grand Rapids, MI) from 1 ng of total RNA using Takara SMART-Seq Stranded Kit (Takara Bio USA, Mountain View, CA) per the manufacturer’s protocol. In brief, RNA was sheared to 300-400 bp, after which dsDNA was generated using a template-switching mechanism, and unique dual-indexed adapters were added to each sample. Ribosomal cDNA was degraded by scZapR and scrRNA probes and libraries amplified with 12 cycles of PCR. The quality and quantity of the finished libraries were assessed using a combination of Agilent DNA High Sensitivity chip (Agilent, Santa Clara, CA), QuantiFluor® dsDNA System (Promega Corp., Madison, WI), and Kapa Illumina Library Quantification qPCR assays (Kapa Biosystems). Individually indexed libraries were pooled and 100 bp, single -end sequencing was performed on an Illumina NovaSeq6000 sequencer (Illumina Inc., San Diego, CA) to return a minimum read depth of 30M read pairs per library. Base calling was done by Illumina RTA3, and the output of NCS was demultiplexed and converted to FastQ format with Illumina Bcl2fastq v1.9.0. To increase read depth and power, the same libraries were re-sequenced to achieve a minimum aligned read depth of 30M read pairs per sample. Technical replicates were combined after generating read counts using the ‘CollapseReplicates’ in DESeq2^75^. FastQ files for both technical replicates will be available at the NCBI Gene Expression Omnibus upon publication.

### RNA seq analysis

RNA-sequencing (RNA-seq) analysis from FastQ to count files was supported through computational resources provided by the Institute for Cyber-Enabled Research at Michigan State University. Quality control of RNA-seq data was performed using FastQC v0.11.7 (Babraham Bioinformatics) and compiled using MultiQC v1.7.^76^ Low-quality bases and adapters were trimmed using TrimGalore v0.6.5 (Babraham Bioinformatics). Reads were aligned to the GRCm38 (mm10) mouse genome using STAR v2.7.3a, and reads were counted using HTSeq v0.11.2.

Count visualization, normalization, and differential expression analysis were performed in R v4.0.5 and utilized packages dplyr v.1.0.5, tidyverse v1.3.0, and readr v1.4.0 to organize data. Low-expression read counts were filtered from DESeq2 normalized counts tables (median of ratios method) using zFPKM v1.12.0 and excluded from the analysis. BioMart was used to generate chromosome X and Y annotations to remove these genes from analysis for control animal comparisons between males and females. Principal component analyses (PCA) and volcano plots were generated in DESeq2 v1.30.1 and visualized using ggplot2 v3.3.3 with ggforce v0.3.3 and ggrepel 0.9.1. Hierarchical clustering heatmaps were generated in DESeq2 v1.30.1 and visualized using pheatmap v1.0.12. Count normalization and differential expression were performed in DESeq2 v1.30.1 where modeling adjusted for RIN and results shrunk using ashr v2.2-47. Gene symbol and other identifying information were determined from Ensembl ID using AnnotationDbi v1.52.0 and org.Mm.eg.db v3.12.0. Knitr v.1.31 was used to export R package citations. Pathway analyses were performed using Gene Set Enrichment Analysis (GSEA) v4.1.0 and Ingenuity Pathway Analysis (IPA) (QIAGEN Inc., https://www.qiagenbioinformatics.com/products/ingenuity-pathway-analysis). Scripts and code utilized in this analysis are available upon request.

### Interactome Analysis of colon-innervating DRG neurons and NMS enteric glia

Published transcriptomic data from sensory neurons was obtained from a single-cell dataset of colon-innervating DRG neurons (GSE102962)^34^. The trinarization score^77^ was calculated for all genes across all neuron populations in the single-cell dataset, and genes with a score of > 0.95, indicating a high likelihood of expression in the neurons, were included in the analysis. Each interactome was generated to show the ligands in one cell type and corresponding receptors in another. Interactomes were generated to show ligands and receptors in the glia that were differentially expressed following NMS in each sex (adjusted p-value < 0.05) and the corresponding receptors and ligands in DRG neurons.

### Immunohistochemistry

Preparations of mouse colonic myenteric plexus prepared from segments of wild-type or NMS intestines preserved in 4% PFA 2 hours at room temperature and were processed for whole-mount immunohistochemistry as previously described^18,78^. LMMP whole mounts were rinsed three times for 10 minutes each with 0.1% Triton X-100 in PBS, followed by 45 minutes incubation in blocking solution (4% normal donkey serum, 0.4% Triton X-100, and 1% bovine serum albumin). Preparations were incubated with primary antibodies overnight at 4°C and secondary antibodies for 2 hours at room temperature before mounting in Prolong Diamond antifade (see **Table S6**).

### Fluorescence in situ hybridization (RNAscope)

LMMP tissue preparations from the distal colon were collected from wild-type and NMS mice at 20 weeks old. Samples were generated following the fixation and dissection procedures described above. RNAscope was performed using the Advanced Cell Diagnostics (ACD) RNAscope 2.5 HD HD Assay-RED reported elsewhere^79^. Tissues were dehydrated and rehydrated by a serial ethanol gradient (25%, 50%, 75%, and 100% in PBS with 0.1% Triton X-100) before H_2_O_2_ treatment. Tissues were then digested with Protease III for 45 minutes and incubated with an RNAscope probe (**Table S6**). Tissues were washed three times for 5 minutes between each step with PBS (before probe incubation) or with RNAscope™ wash buffer (between amplification steps). Labeling was confirmed with species-specific positive and negative controls. Immunohistochemistry was performed following the completion of the RNAscope protocol as described above.

### Imaging

Immunohistochemistry and RNAscope fluorescent labeling were evaluated using the 20 X (0.75 numerical aperture, Plan Fluor; Nikon) of an upright epifluorescence microscope (Nikon Eclipse Ni) with a Retiga 2000R camera (Qimaging) controlled by Qcapture Pro 7.0 (Qimaging) software. Representative images were acquired through the Zeiss LSM 880 NLO confocal system (Zeiss) using Zen Black software and a 20x objective (0.8 numerical aperture, Plan ApoChromat; Zeiss).

### Ca^2+^ imaging

Live whole-mounts of colon myenteric plexus from either *Wnt1^Cre;GCaMP5g-tdT^::GFAP-hM3Dq* from control or NMS mice were prepared for Ca^2+^ imaging as previously described^14^. Whole-mount circular muscle-myenteric plexus (CMMP) preparations were micro-dissected from mouse distal colons. The preparations were continuously perfused with fresh, prewarmed (37°C) Kreb’s buffer consisting of 121 mM NaCl, 5.9 mM KCl, 2.5 mM CaCl_2_, 1.2 mM MgCl_2_, 1.2 mM NaH_2_PO_4_, 10 mM HEPES, 21.2 mM NaHCO3, 1 mM pyruvic acid, and 8 mM glucose (pH adjusted to 7.4 with NaOH) at a flow rate of 2-3 mL.min^-1^. CNO at 10 μM, was applied locally to the ganglion surface to provoke a response. To do this, glass micropipettes were fabricated with a pipette puller (P-87 Flaming-Brown Micropipette Puller, Sutter Instruments Corporation) and capillary glasses (0.85 mm ID/1.25 mm OD, end-opening 60-80 µm of diameter) and back-filled with drug dissolved in Kreb’s Buffer. Drugs were then applied using very gentle positive pressure applied with a 5 mL syringe connected to a pipette holder. This approach delivered microliter volumes of drugs and affected only the ganglion of interest. A previous publication^27^ confirmed that the shear fluid stress associated with drug application did not activate neurons or glia under these conditions. For tetrodotoxin (TTX) treatment, preparations were incubated for 2 minutes with TTX [300 nM], then Kreb’s buffer flow was established to rinse for 1 minute, and CNO stimulation was applied. For EP3 receptor antagonist experiments, an initial stimulation with CNO was done. Then, preparations were incubated with an EP3 receptor antagonist L-798,106 [10 µM] and stimulated with an identical CNO stimulation, allowing us to have the response before and after EP3 receptor antagonism. Images were acquired every second through a 20x plus water-immersion objective lens (Olympus XLUMPLFLN20xW, 1.0 numerical aperture). A DG4 Xenon light source (Sutter Instrument) provided illumination for fluorescence imaging.

### In vivo assessment of visceral sensitivity

Visceral sensitivity was measured as previously described by non-invasive assessment of visceromotor responses (VMRs) to colorectal distensions (CRDs)^18,80^. Intracolonic pressure was measured by a miniaturized pressure transducer catheter (SPR-524, Mikro-Tip catheter, Millar Instruments) equipped with a custom-made plastic balloon. Mice were trained to restrainer for 3 hours one day before the experiment. Mice received an enema of saline solution to remove any pellets in the distal colon. Mice were briefly anesthetized with isoflurane while the pressure transducer balloon was inserted into the colorectum. Mice were then acclimated to the restrainer for 30 minutes before starting the CRD procedure. Graded phasic distensions (20, 40, 60, and 80 mmHg, 2 times each, 20 seconds duration, 3 minutes interstimulus interval) were delivered to the balloon by a Barostat (G&J Electronics) and VMR recordings were acquired using LabChart 8 (AD instruments, CO, USA). For the experiment with AM251, after anesthesia with isoflurane the animal received an intraperitoneal injection of AM251 [1 mg.kg^-1^] and then were acclimated for 30 minutes before starting the CRD procedure.

### In vivo assessment of gastrointestinal transit time

Mice were weighed on the day of the experiments. Animals received 200 µL by gavage of a solution containing 60 mg.mL^-1^ of Carmine red in 0.5% Methylcellulose and received an intraperitoneal injection of CNO [0.25 mg.kg^-1^]. Then, mice were housed in individual cages with *ad libitum* food and water, without bedding, in a behavior room at 21°C, Zeitgeber +2. The fecal pellets were harvested until a red fecal pellet was expulsed. The gastrointestinal transit time was assessed by the difference in time between the gavage and expulsion of the first red feces. Fecal water content corresponded to the weight difference between wet and dry pellets.

### Ex vivo assessment of intestinal contractility

We performed colon and ileum contractility studies as previously described^14^. Briefly, 1 cm strips of fresh tissue were placed in organ chambers (Radnoti, CA, USA) containing 30 mL of Krebs solution at 37°C, continuously bubbled with 95% O_2_ and 5% CO_2_. Isometric contractions were recorded under 1 g passive tension with a force transducer (Grass Instruments, Quincy, MA) coupled with a PowerLab 8/35 (AD instruments) and the LabChart v8.1.5 data analysis software.

Tissues were equilibrated for 30 minutes and then stimulated by bethanechol [10 µM] and CNO [10 µM] to activate enteric glia or electrical field stimulation (EFS) using a RADSTIM electrical stimulator (Radnoti) to activate enteric neurons (train duration 3 s, frequency 20 Hz, pulse duration 400 µs, amplitude 20 V). The induced contractile response was analyzed in basal condition, in the presence of TTX [300 nM], in the presence of L-161,982 [10 mM], an EP3 receptor antagonist, or L-798,106 [10 mM], an EP4 receptor antagonist.

### Analysis and Statistics

Male and female mice were analyzed separately. Whole animals were considered individual n’s for immunoassay, immunohistochemistry, *in situ* hybridization, and *in vivo* analysis. Individual cellular responses from multiple mice represented n’s for Ca^2+^ imaging. Immunolabeling data were analyzed in FIJI (National Institutes of Health) by manually counting positively labeled cells and normalizing them by area. Ca^2+^ recordings were motion-corrected using the FIJI Package StackReg (Philippe Thévenaz, EPFL Switzerland). Ca^2+^ imaging traces represent the change in fluorescence over time of an individual glial cell. Individual glial cells were identified by tdTomato expression and morphology^81^. Calcium recordings were processed by FIJI with the following parameters: Smooth, Substrack background (rolling: 150), and divided by the mean fluorescence of frames 1 to 15 (considered as baseline frame). ΔF/F_0_ from each ROI over time was extracted with Mesmerize software^82^. Raw Ca^2+^ traces were then analyzed with OriginPro 2023b (OriginLab©) to measure and detect peaks with the following parameters: custom baseline =1, peak detected as an increase of at least 20% of the highest peak, and an increase of 25% from baseline, and evaluation of peak duration was assessed using fitting peak features by gaussian extrapolation with fix value for center/height of peaks. Raw VMR traces were processed by running the SmoothSec and root mean square functions in LabChart 8 software to filter phasic colonic contractions^18^. Following filtering, the integral from the response at each distension pressure and the baseline mean from 20 seconds prior to the distension were obtained. Responses were considered significant if they were at least 2 SDV above the baseline mean. Ex vivo assessment of motility results was analyzed with LabChart8 software. The baseline was measured by the maximum value 10 seconds before the stimulation, and the maximum response was assessed by the maximum value observed in the 5 minutes following stimulation. A ratio of maximum response on the maximum value prior to stimulation was calculated to assess the effect of each drug. Statistical analysis of RNA-seq was performed by DESeq2, where adjusted p-value < 0.05 was considered significant. adjusted p-value < 0.1 and |log2fold-change| > 1 were used to perform IPA.

Data were analyzed using Students’ t-test, one-way or two-way ANOVA using GraphPad 10 software (Prism) and are presented as mean ± standard error of the mean (SEM). Outliers were excluded using the outliers ROUT test (Q=1%) from GraphPad.

### Chemicals and Reagents

Chemicals and reagents are listed in **Table S5**.

## Author Contributions

Study concept and design (JG, CD, BDG), acquisition of data (JG, CD, KM, WMS, JM), analysis and interpretation of data (JG, CD, KM, WMS, JM, RN, TJP, BDG), drafting of the manuscript (JG), critical revision of the manuscript (JG, CD, KM, AM, RN, TJP, BDG), statistical analysis (JG, CD, KM, RN), obtained funding (TJP, BDG).

## SUPPLEMENTAL MATERIALS

**Fig. S1.**
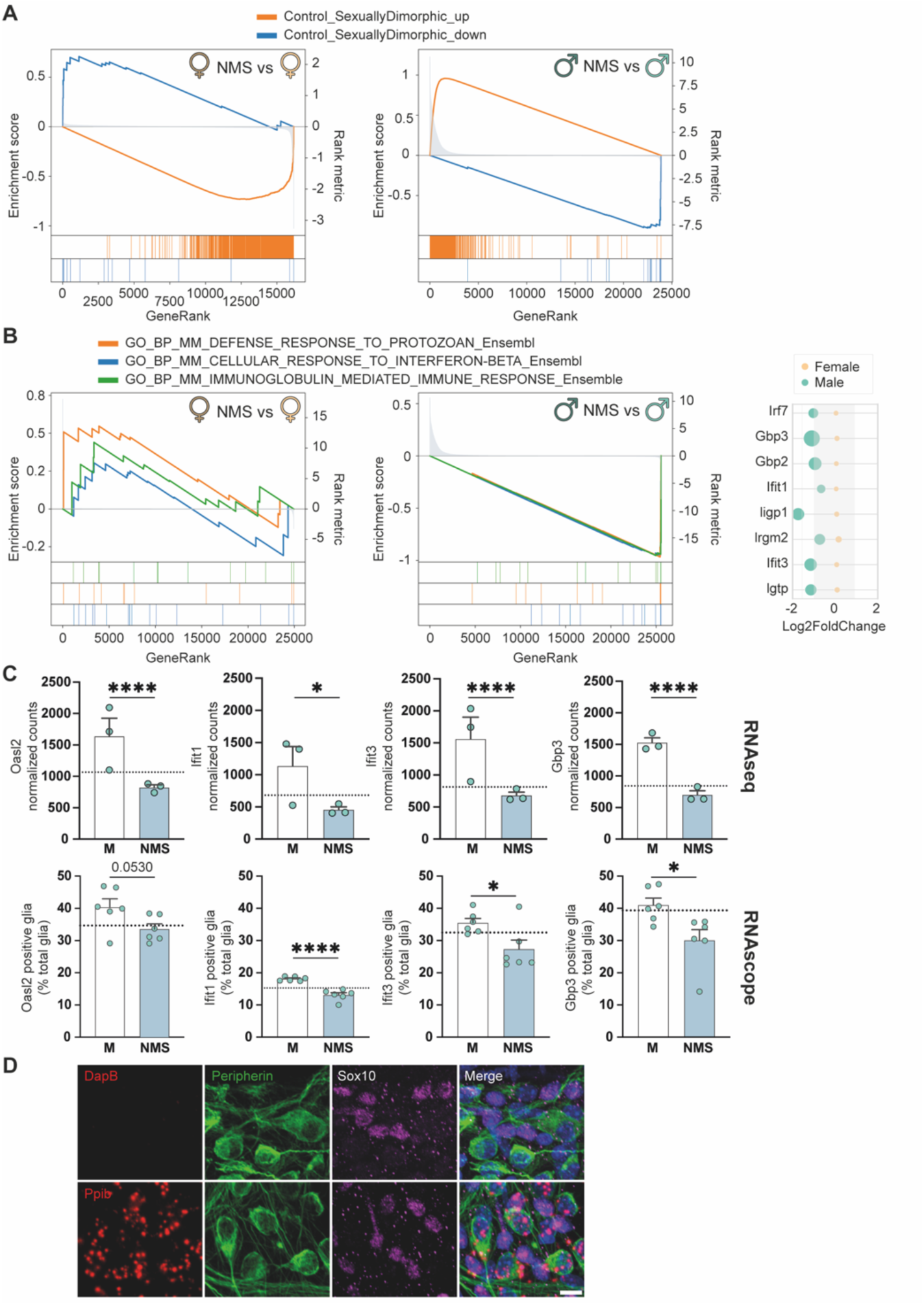
Early life stress induces a male glial gene expression profile similar to females. (**A**) Gene set enrichment analysis of sexually dimorphic genes, identified as differentially expressed genes between control males and females with a |fold-change| ≥ 4, for NMS male and female enteric glia cells DEGs. (**B**) Gene set enrichment analysis of top inflammatory signaling gene sets and gene-wise differential expression from this gene set in males (green) and females (yellow) following NMS. (**C**) (Top) RNA-seq results of IFN signaling molecules *Oasl2*, *Ifit1*, *Ifit3* and *Gbp3*, and (Bottom) quantification of positive enteric glia for RNAscope labeling targeting *Oasl2*, *Ifit1*, *Ifit3* and *Gbp3* in control (white) and NMS (blue) males. Graph bars represent mean + SEM, each individual dot represents the value of one animal, and the dashed line represents the expression level of control females. Values for RNAscope groups were analyzed via student t-test, while RNAseq normalized count represent adjust p-value (* p< 0.05, ** p< 0.01, *** p< 0.001, **** p< 0.0001, n = 6-7 animals per group) RNAscope quantification represents an average of at least 500 enteric glial cells per animal. (**D**) RNA scope negative control (*DapB*) shows no labeling in the myenteric plexus while positive control (*Ppib*) shows labeling. Scale = 10 µm.

**Fig. S2.**
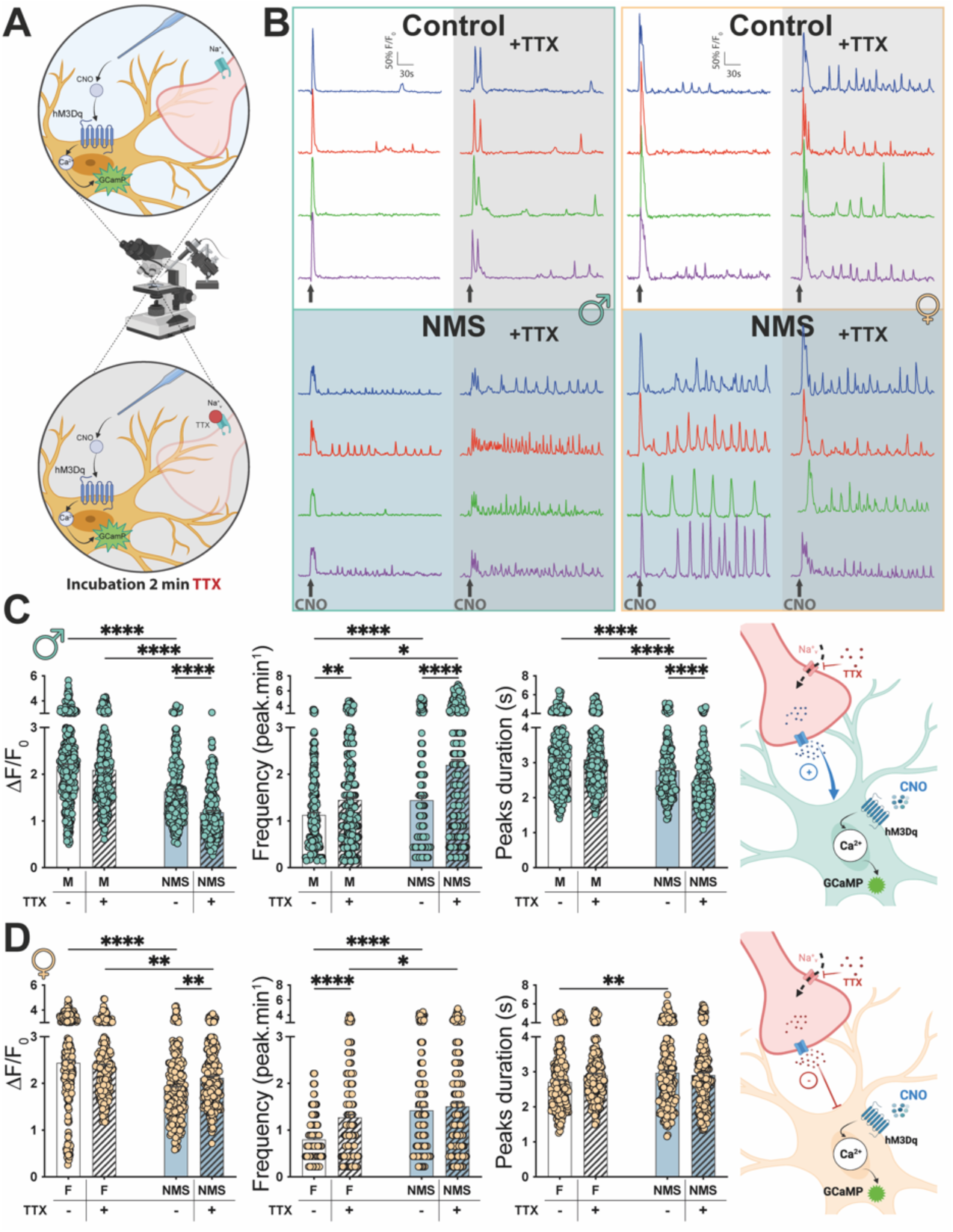
Neuronal input on enteric glia communication is sexually altered after early life stress. (**A**) Schematic methodology of the experiment. (**B**) Representative traces reported as ΔF/F_0_ over time of individual cell’s calcium response following a CNO puffing stimulation (arrow) in control males (Top left) and females (Top right), NMS males (Bottom left) and females (Bottom right) in the absence (white) or in the presence of TTX (grey). (**C-D**) Quantification of maximal peak ΔF/F_0_ response, frequency of response, and maximal peak response duration following CNO stimulation in control (white) and NMS (blue) (**C**) males (green dots) and (**D**) females (yellow dots) animals. (**E**) Schematic representation of proposed mechanisms altered after NMS in male (left) and female (right) enteric glia. Graph bars represent mean + SEM, and each individual dot represents the value of one glial cell. Values for each group were analyzed via two-way ANOVA followed by Bonferroni’s post hoc analysis (* p< 0.05, ** p< 0.01, *** p< 0.001, **** p< 0.0001, n = 230-390 cells from 3-5 animals per group (from at least 3 different ganglia per animal)).

**Fig. S3.**
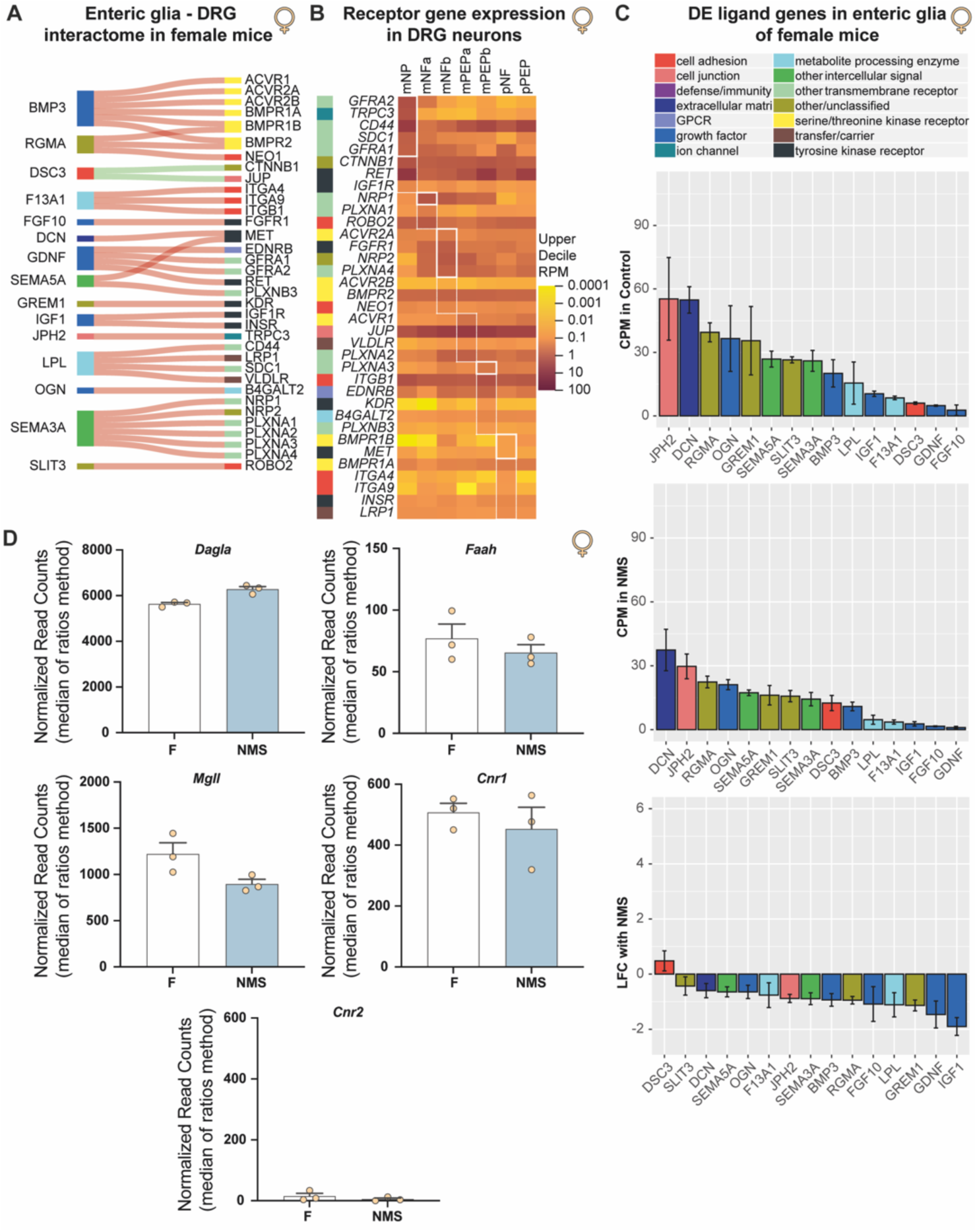
Interaction of female enteric glia with dorsal root ganglia is affected by early life stress, associated with alteration of visceral sensitivity. (**A**) Ligand genes in female enteric glia that were differentially expressed with NMS (adj. p-value <0.05) and their receptor pairs that were expressed in DRG neurons. Green flows indicate upregulation of the ligand gene, and red flows indicate downregulation following NMS. (**B**) Heatmap of receptor gene expression in different populations of colon-innervating DRG sensory neurons; the thick white box indicates enrichment of the gene in the corresponding neuronal population, and the thin dotted box indicates the population with the highest expression of a nonspecific gene. (**C**) Expression of the ligand genes in control and NMS enteric glia (error bars indicate standard deviation) and log2 fold-change with NMS (error bars indicate standard error). (**D**) RNA seq results of *Dagla, Faah, Mgll, Cnr1 and Cnr2* normalized-counts in control (white) and NMS (blue) females (n=3 per group).

**Table S1.** Measurement of in vivo motility and comparison between sexes for gastrointestinal transit time and water fecal content. Values for each group were analyzed via student t-test or Mann-Whitney test; bold p-values are significantly different.

| Conditions |  | Gastrointestinal total transit time (min) |  | Statistic |
| --- | --- | --- | --- | --- |
| Treatment | Genotype | Average Male (± SEM) | Average Female (± SEM) | p-value |
| Control | GFAP-hM3Dq (-) | 261.7 (± 21.97) | 289.4 (± 30.57) | $p = 0.4745$ |
| Control | GFAP-hM3Dq (-) | 265.6 (± 36.91) | 280.6 (± 33.38) | $p = 0.7661$ |
| NMS | GFAP-hM3Dq (+) | 382.1 (± 26.93) | 405.5 (± 39.70) | $p = 0.6255$ |
| NMS | GFAP-hM3Dq (+) | 283.3 (± 38.06) | 423.9 (± 51.77) | **$p = 0.0476$** |

| Conditions |  | H2O in feces (%) |  | Statistic |
| --- | --- | --- | --- | --- |
| Treatment | Genotype | Average Male (± SEM) | Average Female (± SEM) | p-value |
| Control | GFAP-hM3Dq (-) | 50.38 (± 0.75) | 51.23 (± 1.51) | $p = 0.5758$ |
| Control | GFAP-hM3Dq (-) | 54.56 (± 1.04) | 53.95 (± 1.32) | $p = 0.7186$ |
| NMS | GFAP-hM3Dq (+) | 51.44 (± 1.58) | 49.49 (± 1.163) | $p = 0.0950$ |
| NMS | GFAP-hM3Dq (+) | 54.88 (± 1.21) | 57.45 (± 2.356) | $p > 0.9999$ |

**Table S2.** Isolated organ bath chambers ex vivo motility comparison between control and NMS in distal colon and distal ileum. Values for each group were analyzed via student t-test or Mann-Whitney test; bold p-values are significantly different.

Colon
| Conditions | Max value (% of baseline) |  | Statistic |  |
| --- | --- | --- | --- | --- |
| Stimulation | Average control (± SEM) | Average NMS (± SEM) | n (Control - NMS) | p-value |
| Bethanechol | 235.5 (± 17.55) | 197.0 (± 11.07) | 26 - 29 | $p = 0.1411$ |
| EFS | 120.5 (± 4.70) | 131.1 (± 9.12) | 21 - 22 | $p = 0.8709$ |
| CNO | 252.2 (± 14.01) | 208.2 (± 11.33) | 26 - 29 | $p = \mathbf{0.0170}$ |
| TTX +EFS | 99.0 (± 0.84) | 99.8 (± 0.44) | 21 - 22 | $p = 0.4160$ |
| TTX +CNO | 115.6 (± 1.34) | 113.2 (± 1.23) | 26 - 25 | $p = 0.2035$ |
| EP4+CNO | 112.4 (± 1.56) | 110.9 (± 1.82) | 24 - 23 | $p = 0.5406$ |
| EP3+CNO | 113.5 (± 1.68) | 110.1 (± 2.14) | 25 - 27 | $p = 0.2189$ |

Ileum
| Conditions | Max value (% of baseline) |  | Statistic |  |
| --- | --- | --- | --- | --- |
| Stimulation | Average control (± SEM) | Average NMS (± SEM) | n (Control - NMS) | p-value |
| Bethanechol | 157.1 (± 4.32) | 152.6 (± 3.84) | 27 - 29 | $p = 0.4415$ |
| EFS | 113.2 (± 4.34) | 131.3 (± 7.56) | 22 - 22 | $p = 0.0436$ |
| CNO | 216.5 (± 9.73) | 195.0 (± 8.41) | 27 - 29 | $p = 0.0997$ |
| TTX +EFS | 101.3 (± 1.37) | 100.4 (± 1.04) | 22 - 22 | $p = 0.3538$ |
| TTX +CNO | 125.4 (± 2.77) | 119.2 (± 1.69) | 27 - 25 | $p = 0.0679$ |
| EP4+CNO | 123.9 (± 2.24) | 117.5 (± 3.18) | 25 - 22 | $p = \mathbf{0.0330}$ |
| EP3+CNO | 124.2 (± 2.73) | 117.0 (± 2.83) | 25 - 25 | $p = 0.0726$ |

**Table S3.** Isolated organ bath chambers ex vivo motility comparison between males and females in control and NMS distal colon and distal ileum. Values for each group were analyzed via student t-test or Mann-Whitney test; bold p-values are significantly different.

Colon
| Conditions |  | Max value (% of baseline) |  | Statistic |  |
| --- | --- | --- | --- | --- | --- |
| Treatment | Stimulation | Average Male (± SEM) | Average Female (± SEM) | n (M - F) | p-value |
| Control | Bethanechol | 222.9 (± 20.34) | 248.0 (± 29.04) | 13 - 13 | $p = 0.7623$ |
| | EFS | 128.3 (± 7.22) | 111.8 (± 4.83) | 11 - 10 | $p = 0.0782$ |
| | CNO | 260.9 (± 18.87) | 243.6 (± 21.21) | 13 - 13 | $p = 0.5478$ |
| | TTX +EFS | 99.9 (± 1.33) | 98.0 (± 0.95) | 11 - 10 | $p = 0.2849$ |
| | TTX +CNO | 115.1 (± 1.51) | 116 (± 2.27) | 13 - 13 | $p = 0.7282$ |
| | EP4+CNO | 113.5 (± 2.28) | 111.2 (± 2.17) | 12 - 12 | $p = 0.4614$ |
| | EP3+CNO | 111.1 (± 1.70) | 119.8 (± 4.55) | 13 - 13 | $p = 0.2428$ |
| Treatment | Stimulation | Average Male (± SEM) | Average Female (± SEM) | n (M - F) | p-value |
| NMS | Behtanechol | 188.1 (± 12.01) | 203.3 (± 17.03) | 12 - 17 | $p = 0.5069$ |
| | EFS | 123.0 (± 10.52) | 136.7 (± 13.74) | 9 - 13 | $p = 0.4761$ |
| | CNO | 222.4 (± 14.32) | 198.2 (± 16.39) | 13 - 17 | $p = 0.2996$ |
| | TTX +EFS | 99.1 (± 0.61) | 100.2 (± 0.60) | 9 - 13 | $p = 0.2228$ |
| | TTX +CNO | 110.3 (± 1.48) | 115.5 (± 1.67) | 11 - 14 | $p = 0.0351$ |
| | EP4+CNO | 108.9 (± 2.76) | 113.1 (± 2.28) | 12 - 11 | $p = 0.1299$ |
| | EP3+CNO | 108.7 (± 3.51) | 111.2 (± 2.73) | 12 - 15 | $p = 0.3213$ |

Ileum
| Conditions |  | Max value (% of baseline) |  | Statistic |  |
| --- | --- | --- | --- | --- | --- |
| Treatment | Stimulation | Average Male (± SEM) | Average Female (± SEM) | n (M - F) | p-value |
| Control | Bethanechol | 154.3 (± 4.26) | 160.1 (± 7.83) | 14 - 13 | $p = 0.5148$ |
| | EFS | 116.4 (± 7.11) | 109.3 (± 4.40) | 12 - 10 | $p = 0.4266$ |
| | CNO | 219.0 (± 16.56) | 213.7 (± 10.23) | 14 - 13 | $p = 0.7936$ |
| | TTX +EFS | 99.6 (± 1.54) | 103.3 (± 2.32) | 11 - 10 | $p = 0.1916$ |
| | TTX +CNO | 126.7 (± 4.39) | 124.0 (± 3.42) | 14 - 13 | $p = 0.6308$ |
| | EP4+CNO | 120.2 (± 2.12) | 131.5 (± 5.04) | 13 - 13 | $p = 0.0489$ |
| | EP3+CNO | 116.4 (± 1.64) | 127.6 (± 3.91) | 12 - 11 | $p = 0.0130$ |
| Treatment | Stimulation | Average Male (± SEM) | Average Female (± SEM) | n (M - F) | p-value |
| NMS | Bethanechol | 152.6 (± 4.47) | 152.6 (± 5.86) | 12 - 17 | $p = 0.9987$ |
| | EFS | 127.4 (± 7.85) | 134.1 (± 11.81) | 9 - 13 | $p = 0.6755$ |
| | CNO | 213.1 (± 12.94) | 182.3 (± 10.25) | 12 - 17 | $p = 0.0703$ |
| | TTX +EFS | 100.1 (± 1.69) | 100.5 (± 1.36) | 9 - 13 | $p = 0.7401$ |
| | TTX +CNO | 119.8 (± 2.78) | 118.7 (± 2.16) | 11 - 14 | $p = 0.7499$ |
| | EP4+CNO | 117.1 (± 4.37) | 113.8 (± 2.73) | 11 - 10 | $p = 0.5342$ |
| | EP3+CNO | 122.0 (± 3.76) | 113.0 (± 3.88) | 11 - 14 | $p = 0.1145$ |

**Table S4.** Isolated organ bath chambers ex vivo motility comparison by sexes between control and NMS in distal colon and distal ileum. Values for each group were analyzed via student t-test or Mann-Whitney test; bold p-values are significantly different.

Colon
| Conditions |  | Max value (% of baseline) |  | Statistic |  |
| --- | --- | --- | --- | --- | --- |
| Sex | Stimulation | Average Control (± SEM) | Average NMS (± SEM) | n (M - F) | p-value |
| Males | Bethanechol | 222.9 (± 20.34) | 188.1 (± 12.01) | 13 - 12 | $p = 0.3203$ |
| | EFS | 128.3 (± 7.22) | 123.0 (± 10.52) | 11 - 9 | $p = 0.6749$ |
| | CNO | 260.9 (± 18.87) | 222.4 (± 14.32) | 13 - 12 | $p = 0.1229$ |
| | TTX +EFS | 99.9 (± 1.33) | 99.1 (± 0.61) | 11 - 9 | $p = 0.6338$ |
| | TTX +CNO | 115.1 (± 1.51) | 110.3 (± 1.48) | 13 - 11 | **$p = 0.0366$** |
| | EP4+CNO | 113.5 (± 2.28) | 108.9 (± 2.76) | 12 - 12 | $p = 0.2081$ |
| | EP3+CNO | 111.1 (± 1.70) | 108.7 (± 3.51) | 13 - 12 | $p = 0.2925$ |
| Sex | Stimulation | Average Control (± SEM) | Average NMS (± SEM) | n (M - F) | p-value |
| Females | Bethanechol | 248.0 (± 29.04) | 203.3 (± 17.03) | 13 - 17 | $p = 0.1730$ |
| | EFS | 111.8 (± 4.83) | 136.7 (± 13.74) | 10 - 13 | $p = 0.1424$ |
| | CNO | 243.6 (± 21.21) | 198.2 (± 16.39) | 13 - 17 | $p = 0.0959$ |
| | TTX +EFS | 98.0 (± 0.95) | 100.2 (± 0.60) | 10 - 13 | $p = 0.0558$ |
| | TTX +CNO | 116 (± 2.27) | 115.5 (± 1.67) | 13 - 14 | $p = 0.8406$ |
| | EP4+CNO | 111.2 (± 2.17) | 113.1 (± 2.28) | 12 - 11 | $p = 0.2081$ |
| | EP3+CNO | 119.8 (± 4.55) | 111.2 (± 2.73) | 13 - 15 | $p = 0.2945$ |

Ileum
| Conditions |  | Max value (% of baseline) |  | Statistic |  |
| --- | --- | --- | --- | --- | --- |
| Sex | Stimulation | Average Control (± SEM) | Average NMS (± SEM) | n (M - F) | p-value |
| Males | Bethanechol | 154.3 (± 4.26) | 152.6 (± 4.47) | 14 - 12 | $p = 0.7885$ |
| | EFS | 116.4 (± 7.11) | 127.4 (± 7.85) | 12 - 9 | $p = 0.3166$ |
| | CNO | 219.0 (± 16.56) | 213.1 (± 12.94) | 14 - 12 | $p = 0.7858$ |
| | TTX +EFS | 99.6 (± 1.54) | 100.1 (± 1.69) | 12 - 9 | $p = 0.8546$ |
| | TTX +CNO | 126.7 (± 4.39) | 119.8 (± 2.78) | 14 - 11 | $p = 0.2288$ |
| | EP4+CNO | 120.2 (± 2.12) | 117.1 (± 4.37) | 13 - 11 | $p = 0.5109$ |
| | EP3+CNO | 116.4 (± 1.64) | 122.0 (± 3.76) | 12 - 11 | $p = 0.1740$ |
| Sex | Stimulation | Average Control (± SEM) | Average NMS (± SEM) | n (M - F) | p-value |
| Females | Bethanechol | 160.1 (± 7.83) | 152.6 (± 5.86) | 13 - 17 | $p = 0.4418$ |
| | EFS | 109.3 (± 4.40) | 134.1 (± 11.81) | 10 - 13 | $p = 0.0930$ |
| | CNO | 213.7 (± 10.23) | 182.3 (± 10.25) | 13 - 17 | **$p = 0.0418$** |
| | TTX +EFS | 103.3 (± 2.32) | 100.5 (± 1.36) | 10 - 13 | $p = 0.1801$ |
| | TTX +CNO | 124.0 (± 3.42) | 118.7 (± 2.16) | 13 - 14 | $p = 0.1999$ |
| | EP4+CNO | 131.5 (± 5.04) | 113.8 (± 2.73) | 13 - 10 | **$p = 0.0097$** |
| | EP3+CNO | 127.6 (± 3.91) | 113.0 (± 3.88) | 11 - 14 | **$p = 0.0156$** |

**Table S5.** Chemical and reagents.

| Name | Abbreviation | Reference | Lot | Supplier |
| --- | --- | --- | --- | --- |
| Carmines Red | / | C1022-25G | MKBQ8453V | Sigma |
| Methylcellulose | / | M0262-100G | SLBK0272V | Sigma |
| Clozapine-N-Oxide | CNO | 25457 | 3661-76A | NIH |
| Tetrodotoxin, Citrate, *Fugu* sp | TTX | 554412 | D00122168 | Calbiochem |
| L-161,982 | EP4 antagonist | 10011565 | 0626865-18 | Cayman |
| L-798,106 | EP3 antagonist | 11129 | 0597696-18 | Cayman |
| AM 251 | CB1 reverse agonist | 1117 | 18C/283876 | Tocris |
| Bethanechol | / | Bethanechol chloride 200mg | R105H0 | USP Reference Standard |
| Prolong Diamond antifade | / | P36970 | 2519438 | Thermofisher |
| Triton X-100 | / | X100-100mL | SLCJ7494 | Sigma |
| Paraformaldehyde | PFA | 158127-500G | STBJ3696 | Sigma |
| Bovine Serum Albumin | BSA | 001-000-162 | 146710 | Jackson ImmunoResearch |
| Normal Donkey Serum | NDS | 017-000-121 | 166955 | Jackson ImmunoResearch |
| Quant-IT™ RiboGreen RNA assay Kit | / | R11490 | 2170256 | Thermofisher |
| Quiagen Rneasy Plus Micro Kit | / | 74004 | 163026573 | Quiagen |
| Protein A/G beads | / | 88803 | RL244662D | Thermofisher |
| DL-Dithiothreitol solution | DTT | 646563-10X 5mL | MKCJ6494 | Sigma |
| Cyclohexamide | / | C7698-1G | SLBG6034V | Sigma |
| Protease inhibitor | / | P8340 | 077M4042V | Sigma |
| RNAse inhibitor | / | N2118 | 546073 | Promega |
| Heparin | / | H3393-250KU | SLBP3385V | Sigma |
| RNAse-free water | / | 10977-015 | 1897011 | Invitrogen |
| Nonidet P-40 | / | 74385-1L | BCBL6384V | Sigma |
| Sodium deoxycholate | / | D6750-100G | SLCC4936 | Sigma |
| MgCl<sub>2</sub> | / | 2444-01 | 123660 | J.T.Baker |
| KCl | / | 746436-500G | SLCH0615 | Sigma |
| Tris (1M) pH 7 - RNAse free | / | AM9850G | 1016113 | Invitrogen |
| DMEM/F-12 | / | 11039-21 | 2425956 ; 2558318 ; 2518212 | Gibco |
| NaCl | / | 746398-2.5KG | SLCL3365 | Sigma |
| CaCl<sub>2</sub> | / | 1332-01 | 116456 | J.T.Baker |
| NaH<sub>2</sub>PO<sub>4</sub> | / | 3818-01 | 134199 | J.T.Baker |
| HEPES | / | H3375-1KG | SLCL5410 | Sigma |
| NaHCO<sub>3</sub> | / | 3506-01 | 23H0961034 | J.T.Baker |
| Sodium pyruvate | / | P5280-100G | SLBP4879V | Sigma |
| Glucose | / | G7021-1KG | SLBZ6903 | Sigma |
| Borosilicate Glass with filament | / | BF150-86-10 | 2601321 | Sutter Instrument |
| RNAscope Wash Buffer Reagents | / | 310091 | 2007978 | Advanced Cell Diagnostic (ACD) |
| RNAscope H<sub>2</sub>O<sub>2</sub> | / | 322330 | 2016785 | ACD |
| RNAscope Protease III | / | 322337 | 2015061 | ACD |
| RNAscope 2.5 HD Detection Reagent - RED | / | 322360 | 2012950 | ACD |

**Table S6.** Antibodies and RNAscope probes.

| Primary antibodies |  |  |  |
| --- | --- | --- | --- |
| Name | Reference | Lot | Suppliers |
| HA | 901514 | B222922 | BioLegend |
| Sox10 | HPA068898-100ul | 000C32918 | Atlas Antibodies |
| DAPI | D8417-1mg | 0000116963 | Sigma |
| Peripherin | sc-377093 | I0821 | Santa Cruz Biotechnology |
| Secondary antibodies |  |  |  |
| Name | Reference | Lot | Suppliers |
| Donkey anti-Rabbit Alexa Fluor 594 nm | 711-585-152 | 146903 | Jackson ImmunoResearch |
| Donkey anti-mouse Alexa Fluor 488 nm IgG2a | 115-545-206 | 156607 | Jackson ImmunoResearch |
| Donkey Anti-Rabbit Alexa Fluor 647 nm | 711-605-152 | 149049 | Jackson ImmunoResearch |
| RNAscope probes |  |  |  |
| Name | Reference | Lot | Suppliers |
| *RGS5* | 430181 | 21165B | ACD |
| *Oasl2* | 534501 | 21054B | ACD |
| *Ltbr* | 401021 | 21034A | ACD |
| *Ifit1* | 500071 | 21166C | ACD |
| *Ifit3* | 508251 | 21145A | ACD |
| *Gbp3* | 541801 | 21167A | ACD |
| *DapB* | 310043 | 200959A | ACD |
| *PpiB* | 313911 | 20114A | ACD |
| *UBC* | 310771 | 21090B | ACD |

